# Engineering the ADDobody protein scaffold for generation of high-avidity ADDomer super-binders

**DOI:** 10.1101/2023.09.09.556966

**Authors:** Dora Buzas, Huan Sun, Christine Toelzer, Sathish K. N. Yadav, Ufuk Borucu, Gunjan Gautam, Kapil Gupta, Josh Bufton, Julien Capin, Richard B. Sessions, Frederic Garzoni, Imre Berger, Christiane Schaffitzel

**Affiliations:** School of Biochemistry, University of Bristol, University Walk, Bristol BS8 1TD, UK; Max Planck Bristol Centre for Minimal Biology, Cantock’s Close, Bristol BS8 1TS, UK; The Institute of Medicinal Plant Development (IMPLAD), No. 151 Malianwa North Road, Haidian District, Beijing 100193 P. R. China; Imophoron Ltd, Science Creates Old Market, Midland Rd, Bristol BS2 0JZ UK; School of Chemistry, University of Bristol, Cantock’s Close, Bristol BS8 1TS, UK

**Keywords:** Protein engineering, alternative scaffold, ADDobody, ribosome display selection, ADDomer nanoparticle super-binder

## Abstract

Adenovirus-derived dodecamer (ADDomer) nanoparticles comprise 60 copies of Adenovirus penton base protein (PBP). ADDomer is thermostable, rendering the storage, transport and deployment of ADDomer-based therapeutics independent of a cold-chain. To expand the scope of ADDomer nanoparticles for new applications, we engineered ADDobodies. ADDobodies represent the crown domain of the PBP, genetically separated from its multimerization domain. We inserted heterologous sequences into hyper-variable loops in the crown domain. The resulting ADDobodies were expressed at high yields in *Escherichia coli,* are monomeric and maintain thermostability. We solved the X-ray structure of an ADDobody prototype validating our design. We demonstrated that ADDobodies can be used to select a specific binder against a target in *in vitro* selection experiments using ribosome display, with an enrichment factor of ∼10^4^-fold in one selection round. We show that ADDobodies can be converted back into ADDomers by genetically reconnecting the selected ADDobody with the PBP multimerization domain from a different species, giving rise to a multivalent nanoparticle, called Chimera, confirmed by a 2.2 Å structure determined by cryogenic electron microscopy (cryo-EM). Chimera comprises 60 binding sites, resulting in ultra-high, picomolar avidity to the target.

## Introduction

High-specificity and high-affinity monoclonal antibodies are vital reagents in modern biomedicine and biotechnology. Phage display and yeast display are commonly used technologies to select antibodies using libraries from immunized animals ^1,2^. Nonetheless, both display technologies are limited in the size of the antibody library due to a necessary transformation and amplification step in cells. The ribosome display *in vitro* selection technology avoids any *in vivo* steps that limit the diversity of the library and thus allows selecting binders from very large libraries (up to ∼10^12^). Therefore, ribosome display is particularly suited for naïve and synthetic libraries with a lower content of potential binders (in contrast to antibody libraries generated from immunized animals) ^3,4^. Moreover, ribosome display allows the selection and evolution of antibodies *in vitro* under user-defined selection conditions, independent of the immunogenicity or toxicity of the antigen, and/or also against highly conserved targets ^5,6^. Importantly, affinities in the picomolar and femtomolar range have been reported for *in vitro* selected antibodies, far exceeding the affinities found in natural antibodies ^7,8^.

Alternative binder scaffolds are based on non-immunoglobulin domains and ideally have the following properties: improved biophysical properties, small size and binding of their target with similar or higher affinity and/or specificity as compared to antibodies. Furthermore, alternative binder scaffolds can be produced with high yields as recombinant proteins in *Escherichia coli* (low cost) ^9^. In order to avoid misfolding, optimize yields and permit expression in the reducing environment of the cytoplasm, disulfide bonds or cysteines are usually avoided in alternative binder scaffolds. Alternative binders have many applications including purification, detection and quantification of their target proteins, facilitating structure elucidation, live imaging, for targeted drug delivery and as protein-based therapeutic ^9–15^.

Here, we introduce ADDobodies as a novel alternative binder scaffold for ribosome display selection. ADDobody is derived from ADDomer (<u>A</u>denovirus-derived <u>do</u>deca<u>mer</u>) which we previously developed as a thermostable scaffold with low intrinsic immunogenicity and demonstrated its use as a novel vaccine nanoparticle ^16,17^. ADDomer is based on the PBP of human adenovirus Ad3 serotype and exhibits intrinsic self-organizational properties in the test tube. In the ADDomer, the PBP monomers spontaneously assembles to form a pentamer, and twelve pentamers organize into a dodecamer (Fig. 1A) characterized by high stability and thermotolerance exceeding 50 °C ^16–18^. The PBP monomer folds into two distinctive domains (Fig. 1A,B); the jelly-roll fold domain which is located in the core of the dodecamer and assumed to be responsible for multimerization, and a solvent-facing exterior domain, called the crown domain ^16^. The crown domain comprises two highly flexible regions, called variable loop (VL) and arginine-glycine-aspartic acid (RGD) motif-containing loop (Fig. 1A). These flexible loop regions were identified previously as versatile insertion sites that can be used for displaying foreign sequences such as immunogenic epitopes for genetically encoded multiepitope display ^16,17,19^. The display of multiple pathogen-derived peptide epitopes on the surface of ADDomer enabled the design of vaccine candidates which could elicit strong specific immunoglobulin responses against Chikungunya virus ^16^, SARS-CoV-2 virus ^17,20^ and Foot-and-Mouth Disease ^21^. Importantly, the high thermostability (>50°C) of ADDomer enables refrigeration-independent distribution and storage ^16^. Also, lyophilization of ADDomers is possible ^22^. ADDomer-based thermostable vaccines or therapeutics, due to thermostability, could be deployed independent of a cold chain, which is a highly significant criterium given that each year ∼50% of vaccine doses are estimated to be wasted due to cold chain disruptions world-wide ^23^.

**Figure 1.**
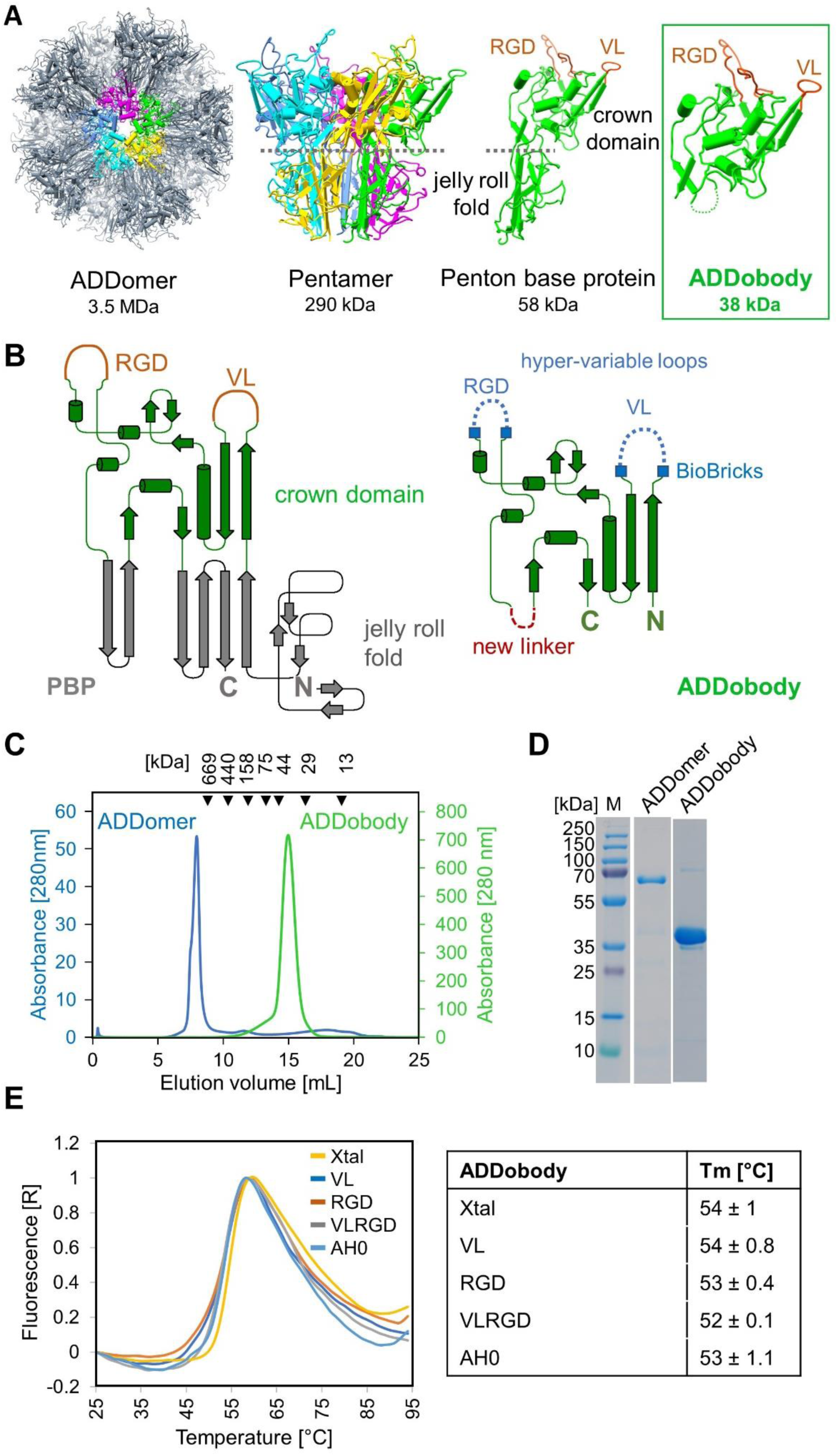
ADDobody protein design. **(A)** ADDomer (left, PDB ID: 6HCR ^16^) assembles spontaneously from twelve pentons (middle left) derived from the adenovirus penton base protein (PBP) (middle right). Each PBP comprises a jelly-roll multimerization domain and a head domain (crown) with flexible VL and RGD loop regions (orange). ADDobody (right) consists of the isolated, engineered crown domain. **(B)** Topology drawing for the PBP (left) as observed in ADDomer and the designed ADDobody (right). In the BioBrick format the transition between the protein scaffold and the hyper-variable loop regions (blue) is marked by restriction site sequences (blue boxes). **(C)** Size exclusion chromatography profile of ADDomer (in the void volume) and monomeric ADDobody which elutes at the expected volume for the size of a ∼38 kDa protein using a Superdex 200 10/300 GL column. **(D)** Coomassie-stained SDS–PAGE showing purified ADDomer/ PBP and ADDobody. Left: Molecular weight marker. **(E)** Thermal unfolding curves of different ADDobody constructs and resulting melting temperatures (Tm) (n=3) are shown for ADDobodies used in this study.

We designed ADDobody, a new scaffold protein based on the crown domain of the penton base (Fig. 1A,B). We show that ADDobodies exhibit the advantages of alternative protein scaffolds outlined above: ADDobodies are single domain, monomeric proteins with comparatively small molecular weight (∼38 kDa), exhibit a highly stable structure without any disulfide bonds and can be produced at high yields using *E. coli.* Of note, ADDobodies retain the thermostability of the ADDomer - our designed ADDobody prototype exhibits a melting temperature (Tm) of 54 °C. We determined two crystal structures of ADDobody at 2.9 Å and 3.2 Å resolution respectively, validating our design. Moreover, we show that ADDobody tolerates insertion of sequences of variable lengths into the hyper-variable loops (VL and RGD), retaining high expression yields and melting temperature of the ADDobody prototype, which is a prerequisite for randomization of the loops to generate a future synthetic ADDobody library. In a proof-of-concept experiment, we demonstrate that ADDobody can be used for *in vitro* selection by ribosome display with an enrichment factor of ∼10^4^-fold per selection round.

Importantly, we show that by genetically fusing ADDobody and the multimerization jelly-roll fold domain, ADDobody can be converted back into an ADDomer nanoparticle, offering 60 binding sites against the target protein used as a bait in ribosome display, resulting in ultra-high avidity. The avidity concept is a useful means to enhance binding to a target, exploited by the immune system for instance by Immunoglobulin G (IgG) antibodies possessing two binding sites, while IgA has four and IgM even ten binding sites for an antigen target. With 60 binding sites, we expect to generate ADDomers that virtually do not release their target. Specifically, we rejoin ADDobody derived from human Adenovirus Ad3 with a jelly-roll fold domain from a Chimpanzee adenovirus, giving rise to a functional chimeric ADDomer nanoparticle, which we call ‘Chimera’. We solve the structure of this Chimera nanoparticle by cryogenic electron microscopy (cryo-EM). Chimera binds its target with ultra-high (picomolar) avidity corroborating our concept that ADDomer-based super-binders can be engineered which bind their target with high efficiency. We anticipate that ADDomer nanoparticles generated by using ADDobodies selected as described, could be highly useful molecular tools, for instance for efficient detection, binding and neutralization of diverse pathogenic or toxic targets.

## Results

### Design and biophysical characterization of ADDobody constructs

We set out to design and produce the crown domain of human Adenovirus Ad3 PBP in isolation, and to determine the biophysical and biochemical properties, and the structure of the resulting protein, which we call ADDobody. For this, the crown domain had to be separated from the jelly-roll fold domain, and the N-zand C-terminal parts reconnected by introducing a linker (Fig. 1A,B). To this end, two beta-strands comprising residues 423-456 were deleted that are part of the jelly-roll fold domain of the Ad3 PBP (Fig. 1B) ^16^. We replaced this part by a flexible 6 amino acid residue linker with the sequence NGDSGN, reconnecting the crown polypeptide chain (Fig. 1B). The VL and RGD loops of the PBP are hyper-variable in length and sequence in nature ^24^ and, as we have shown previously for ADDomer, represent versatile insertion sites for heterologous amino acid sequences seemingly not constrained by length and sequence context ^16^. Therefore, we reasoned that randomized sequences could be introduced also in the VL and RGD loops of ADDobody, facilitating the future generation of a library comprising randomized loops. We chose a ’BioBrick’ format ^25^ to delineate the boundaries of backbone and loops in the ADDobody scaffold. The position of the restriction sites was based on the ADDomer structure (PDB: 6HCR) ^16^ (Fig. 1B).

To validate our design by high-resolutions structural analysis using X-ray crystallography, we created an ADDobody prototype with minimized loops, called ADDobody-Xtal, based on the assumption that long flexible loops could be detrimental to crystal lattice formation. Moreover, we designed exemplar ADDobodies comprising extended loop sequences in the VL loop (ADDobody-VL), in the RGD loop (ADDobody-RGD), in both VL and RGD loops (ADDobody-VLRGD) in BioBrick format. Finally, we designed an ADDobody comprising a viral epitope sequence, AH0, inserted into VL (ADDobody-AH0). The AH0 sequence is part of the receptor binding motif (residues Y473 to Y505) of the Ancestral strain SARS-CoV-2 spike protein ^17^, which mediates binding to the human host cell receptor angiotensin converting enzyme 2 (ACE2)^26^. We have shown previously that the AH0 sequence is recognized and tightly bound by a camelid VHH nanobody, ADAH11 ^17^. The amino acid sequences of the ADDobodies used in this study are provided in Supplementary Table S1.

All ADDobody constructs were successfully expressed in *E. coli* BL21(DE3) with high yields (> 10 mg/L cell culture) within 3 hours at 30 °C. ADDobodies were affinity purified via the His-tag, followed by anion exchange and size exclusion chromatography (SEC) (Fig. 1C,D; Fig. S1). ADDobodies eluted at ∼15 ml from a Superdex200 column as expected for a monomer with a molecular weight of ∼38 kDa (Fig. 1C,D). For comparison, the PBP (∼64 kDa) forms a dodecamer of pentons (the ADDomer) with a size of 3.4 MDa and elutes close to the void volume (∼7 ml) (Fig. 1C).

To determine the stability of our newly designed proteins, purified ADDobodies were analyzed by thermal unfolding using a Thermofluor assay ^27^. The melting temperatures (Tm) for all ADDobody constructs, ranged from 52 °C to 54°C (Fig. 1E) indicating that the ADDobody scaffold adopts a robust fold, irrespective of the individual loop lengths and sequences or the introduction of the linker, none of which seemingly affected thermostability. For comparison, the ADDomer nanoparticle exhibits a Tm of 54 °C ^16^. In summary, we successfully designed an ADDobody scaffold that has excellent expression yields in *E. coli*, maintains thermostability and tolerates diverse amino acid sequence contexts and lengths in its loops, recapitulating favorable characteristics of the ADDomer nanoparticle from which ADDobody is derived.

### Crystal structures of ADDobody

ADDobody with minimized VL and RGD loop sizes (ADDobody-Xtal, Table S1) was crystallized. Crystals grew under two different conditions, one containing 0.1 M zinc acetate, the other devoid of zinc. During X-ray diffraction data collection and structure determination (Table S2), two different crystal forms were observed: In the absence of zinc ions, ADDobody crystals adopted space group P1 and diffracted to a resolution of 3.17 Å.

ADDobody crystals grown in the presence of zinc adopted space group P2_1_2_1_2_1_, and diffracted to a resolution of 2.90 Å. The ADDobody structures from both crystal forms show identical scaffolds while the VL and RGD loops appear flexible (Fig. 2A, Fig. S2). Superimposition of ADDobodies from the crystals with the crown domain from the ADDomer cryo-EM structure revealed a virtually identical fold with a root-mean-square deviation (RMSD) of 1.12 Å (Fig. 2B). Three segments in the ADDobody crystal structures were partially unresolved to varying degrees in the individual chains in the asymmetric units, including VL and RGD loops, and a region close to the C-terminus comprising residues 270 to 305 (Table S1). In the zinc-free ADDobody crystal form, the VL loop could be manually traced through the density in some of the monomers (Fig. S2). The RGD loop region lacked density for about 12-15 residues in all chains of both refined structures indicating conformational disorder (Fig. S2, Fig. 2B).

**Figure 2.**
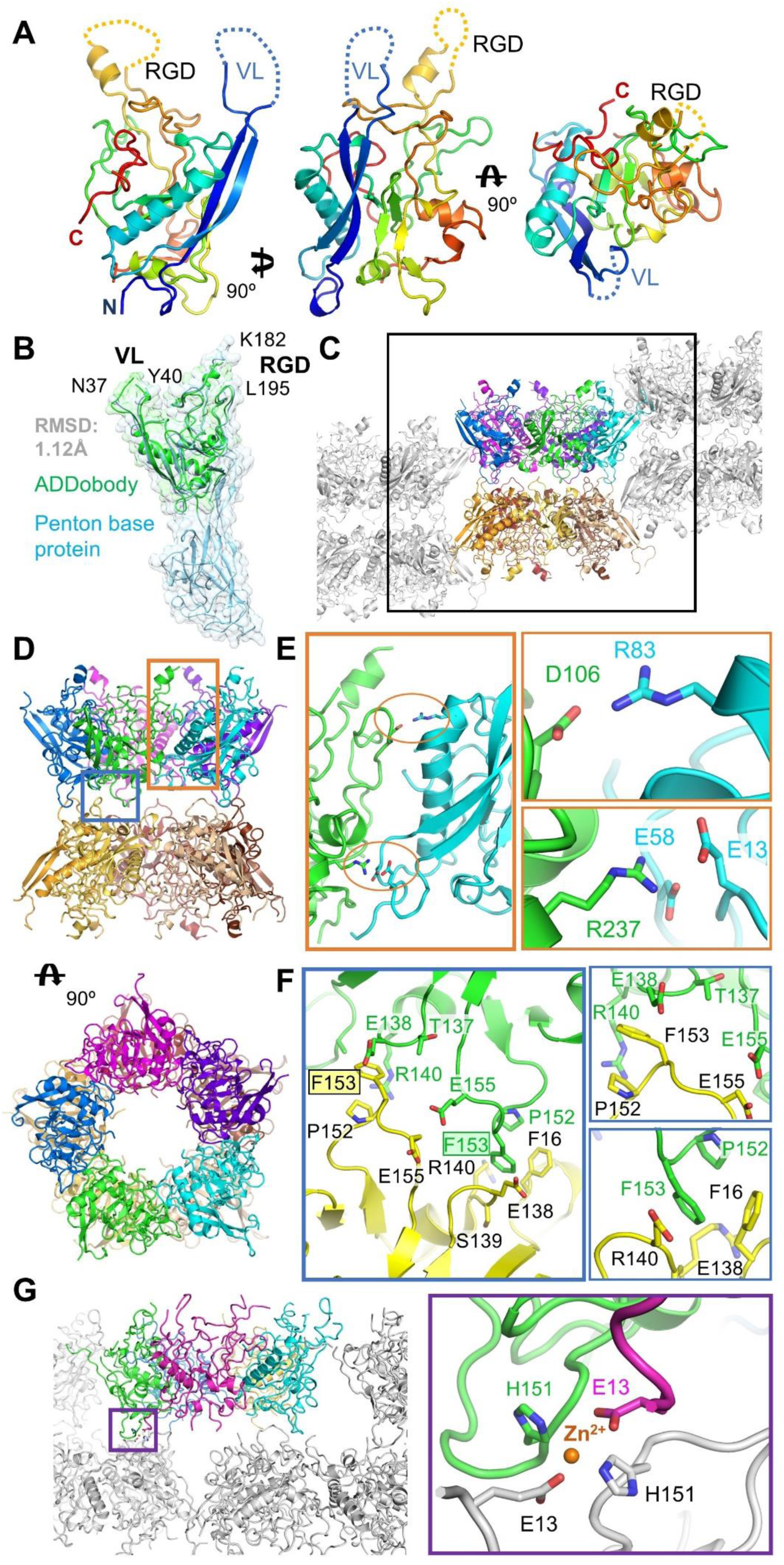
ADDobody crystal structures. **(A)** ADDobody structure in a front, side and top view. The flexible RGD (yellow) and VL (blue) loops are indicated with dashed lines; the N-terminus (blue) and C-terminus (red) are marked. **(B)** Alignment of ADDobody (green) and PBP (cyan, PDB ID: 6HCR ^16^) crown domain with RMSD of 1.12 Å. **(C)** Decameric organization observed in the ADDobody crystal. Two pentons stack together with flexible loop regions facing outwards. A side-view representation is shown with one decamer in color. Black box: unit cell. **(D)** Side and top view of the ADDobody decamer forming a barrel. **(E)** Left: Close-up view of the ADDobody interface within the penton (orange box in panel D). Right: charge-charge interactions stabilizing the interface. **(F)** Left: Close-up view of the interface between ADDobody pentons in the decamer (blue box in panel D). Right: Hydrophobic contacts stabilizing the interface. **(G)** Left: Zn^2+^-containing crystal form showing ADDobody pentons (purple box highlights interface). Right: Zn^2+^ coordination by His151 and Glu13 residues promotes a crystal contact between two symmetry related pentons.

In the zinc-free ADDobody crystal form, twenty ADDobody monomers are found in the asymmetric unit (Fig. 2C). Of note, five ADDobody monomers formed a penton in the crystal reminiscent of the ring-like arrangement in pentons formed by the PBP in the ADDomer (Fig. 2C,D). Two such penton rings in the zinc-free triclinic crystal form a barrel-like decamer where the VL and RGD loops are outward-facing and accessible (Fig. 2C,D). The Coulomb potential indicates a complementary patch of negative and positive charges at the ADDobody surface which contribute to penton ring formation in the crystals (Fig. S3, Fig. 2E). Specifically, D106 forms a charge-charge interaction with R83. E13 and E58 are both in close proximity to R237 (Fig. 2E). The stacking of the two penton rings is mostly mediated by van-der-Waals interactions, with residue F153 playing a central role (Fig. 2F). A cation-п interaction is formed between F153 and R140 of an adjacent chain, and F16 could also interact with F153 of the neighboring ADDobody polypeptide chain (Fig. 2F). The crystallization of ADDobody proteins in the presence of zinc resulted in crystals with a characteristic needle-like shape. In this zinc-containing crystal form, ADDobodies also adopted a pentamer (Fig. 2G), however, the pentameric rings are not stacked on top of each other. Rather, a zinc ion mediates the interaction between pentamer rings. The zinc ion is coordinated by four residues from four different ADDobody chains: H151 donated from one chain, E13 from an adjacent ADDobody chain of the same pentamer and H151 and E13 from the juxtaposed pentameric ring (Fig. 2G, Fig. S4).

Taken together, the crystal structures validated our design, confirming that the crown domain adopts a stable fold with flexible loops. Moreover, the structures imply that multimerization on the level of the pentamer in the ADDomer nanoparticle may not be mediated by the jelly-roll fold domains alone, but that the crown domains also contribute, given that the ADDobodies were found to form pentameric rings in both crystal forms, in spite of being monomeric in solution (Fig. 1C). Metal coordination appears to play an important role in structural organisation - in our previous ADDomer cryo-EM structure, the interaction between crown domain and jelly-roll fold was stabilized by metal binding (likely a zinc ion based on geometry) coordinated by methionine and cysteine residues (PDB ID: 6HCR) ^16^. Here, in our zinc-containing ADDobody crystal structure, we found that the interaction between pentameric ADDobody rings is stabilized by coordinating zinc ions.

### ADDobodies can be used for *in vitro* selection by ribosome display

For a proof-of-concept selection experiment, we cloned two ADDobody variants, ADDobody-AH0 and ADDobody-57. ADDobody-AH0 comprises a viral epitope, AH0, inserted in VL (Table S1). AH0 interacts tightly with a specific target antigen, ADAH11 ^17^. ADDobody-57, in contrast, contains unrelated sequences in VL and RGD loops, and does not bind ADAH11. The constructs were converted into the ribosome display format by removing stop codons and adding a C-terminal spacer sequence derived from TonB fused in frame to the ADDobody-coding sequences ^28^. In addition, a T7 promoter and a ribosome-binding site (Shine Dalgarno sequence, SDA) were introduced ^28^ (Fig. 3A). Importantly, the AH0 encoding DNA sequence comprises a unique restriction site for restriction enzyme *Pst*I. Therefore, *Pst*I restriction digestion can be used to discriminate the PCR product encoding ADDobody-AH0 from the ADDobody-57 encoding construct which remains uncut (Fig. 3A). In agarose gel electrophoresis, the PCR fragment representing uncut ADDobody runs at ∼1620 bp while the *Pst*I-digested AH0 construct gives rise to two bands at ∼1400 bp and 242 bp (Fig. 3A).

**Figure 3.**
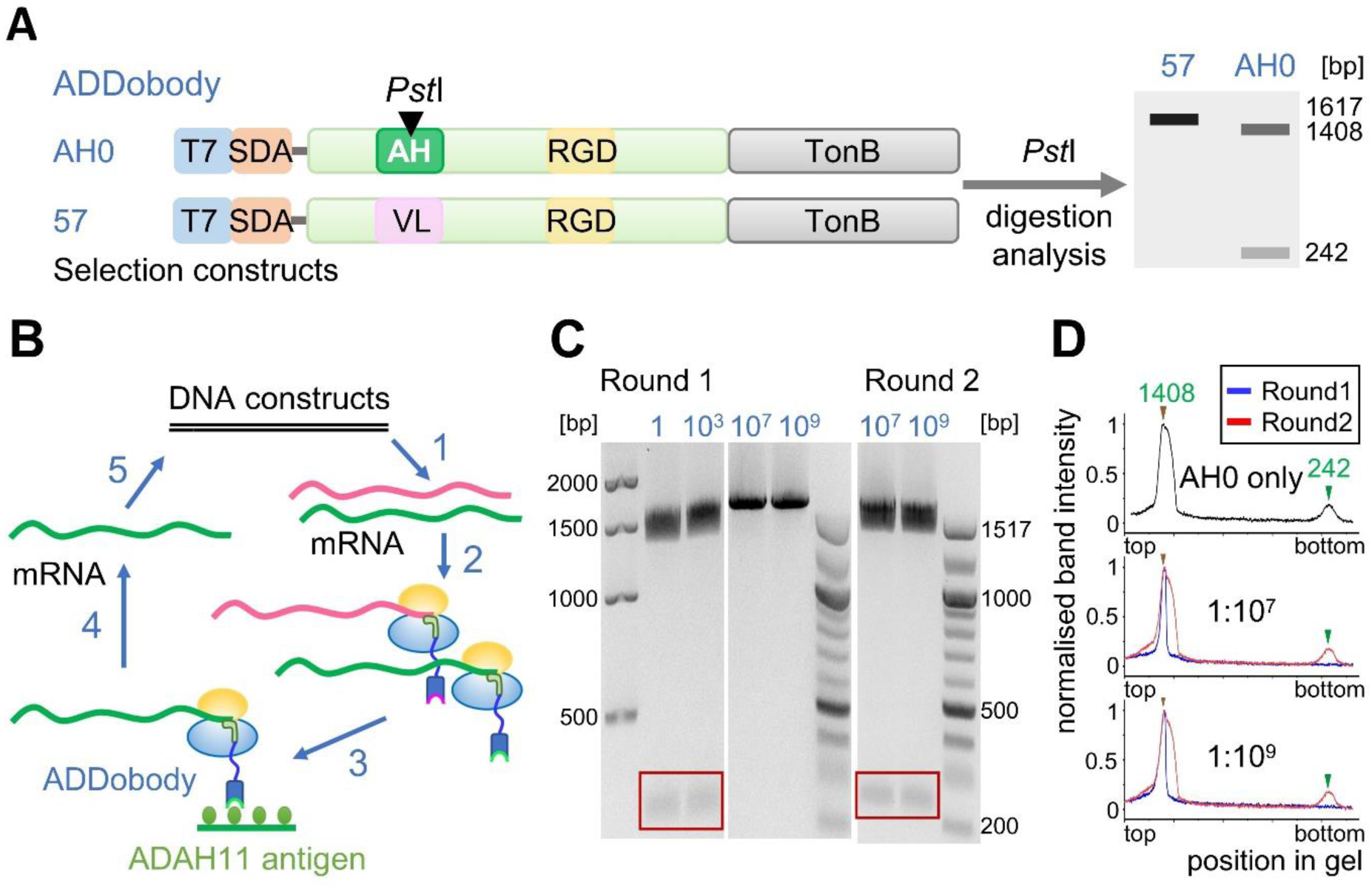
Proof-of-concept ribosome display selection using the ADDobody scaffold. **(A)** Schematic illustration of the ADDobody-encoding gene constructs used for ribosome display. Top: ADDobody-AH0 can bind ADAH11 and its encoding gene has a unique restriction site (*Pst*I). Bottom: The non-binding ADDobody-57. Right: *Pst*I restriction digest of these constructs yields two gene fragments (1408 bp and 242 bp) for ADDobody-AH0, but ADDobody-57 remains uncleaved. **(B)** Schematic illustration of ADDobody ribosome display selection. Gene constructs encoding ADDobodies AH0 and 57 are transcribed *in vitro* (1). mRNA encoding ADDobody-AH0 is mixed with mRNA for ADDobody-57 in defined ratios (up to 1:10^9^) for *in vitro* translation (2). Biotinylated ADAH11 is the bait for selections. Non-binding mRNA-ribosome-ADDobody complexes are washed away (3). The mRNA is eluted (4), reverse transcribed and PCR amplified (5). **(C)** Agarose gel of *Pst*I-digested PCR product after one and two rounds of ribosome display, monitoring enrichment of AH0 binders. Dilutions of AH0 into 57 are indicated in blue. The 242 bp fragment resulting from digestion of ADDobody-AH0 DNA is highlighted (red box). **(D)** Quantification of *Pst*I digestion using DNA agarose gels showing that two rounds of ribosome display (red line) enrich ADDobody-AH0 from 1:10^7^ and 1:10^9^ dilutions. Upper panel: positive control (AH0 only, black line).

Ribosome display *in vitro* selections were performed using biotinylated ADAH11 as the bait (Fig. 3B, Fig. S5). When only the ADDobody-AH0 encoding construct was used for ribosome display selection, the recovered PCR product was completely digested by *Pst*I enzyme (Fig. 3C,D). This confirms that the PCR product encodes ADDobody-AH0 and that ADDobody-AH0 binds its ADAH11 target. Furthermore, this experiment suggests that ADDobody-AH0 is properly displayed by the nascent mRNA-ribosome complexes tethered to the ADDobody due to the deleted stop codon, because the corresponding mRNA encoding ADDobody-AH0 could be efficiently recovered. To test the specificity of binding and the enrichment efficiency of ribosome display selections using ADDobodies, mRNA encoding ADDobody-AH0 was diluted in mRNA encoding ADDobody-57 at a ratio of 1:1, 1:10^3^, 1:10^5^, 1:10^7^, and 1:10^9^. These mRNA mixtures were then used for ribosome display selection experiments (Fig. 3B). We asked whether ADDobody-AH0 could be recovered from this dilution based on ribosome display selection against its target antigen, ADAH11. After the first round of ribosome display selection, ADDobody-AH0 could be detected in the DNA pools of the 1:1 and 1:10^3^ dilutions (Fig. 3C, Fig. S6) as evidenced by the presence of bands of ∼1,400 bp and 242 bp. In contrast, dilutions 1:10^5^, 1:10^7^, and 1:10^9^ resulted in PCR products that could not be digested by *Pst*I (Fig. 3C, Fig. S6), indicating the prevalence of ADDobody-57. These PCR pools were subjected to a second round of ribosome display selection. The resulting PCR pools after round 2 were again *Pst*I digested. All experiments showed at least a partial digestion of the full-size product (bands at ∼1,600 bp, ∼1,400 bp and 242 bp) evidencing co-existence of ADDobody-AH0 and ADDobody-57 (Fig. 3C,D, Fig. S6). Based on these experiments with the 1:10^7^ and 1:10^9^ dilutions of AH0 (Fig. 3D), we estimated an enrichment factor of ∼10,000 for ADDobody-AH0 per ribosome display selection round against its cognate target antigen ADAH11.

To summarize, our ribosome display experiments demonstrate that ADDobody can be properly displayed on the ribosome suitable for *in vitro* selection. The mRNA-ribosome-ADDobody complexes can be used for selection against a cognate target, and ADDobodies binding to the target can be efficiently enriched over non-binding ADDobodies in a ribosome display selection experiment. This is a prerequisite for future selections of ADDobodies from a synthetic library with randomized VL and RGD loop sequences which can conceivably encompass up to 10^12^ members ^29^.

### Design and cryo-EM structure of Chimera super-binder

To test our concept that a selected ADDobody can be reconnected with a jelly-roll fold multimerization domain to yield ultra-high affinity ADDomer-based super-binders, we set out to graft ADDobody-AH0, derived from human Adenovirus Ad3, onto the jelly-roll fold of the Chimpanzee adenovirus serotype Y25 PBP (Fig. 4A) to create a chimeric ADDomer, called Chimera, displaying 60 copies of AH0. This Chimera AH0 nanoparticle was produced using baculovirus/insect cell expression and purified by SEC and IEX to homogeneity (Fig. S7). We confirmed ADDobody-AH0 binding to ADAH11 by SEC and surface plasmon resonance (SPR) (Fig. 4B, Fig. S7B,C). In SEC, a peak was observed at ∼14 ml with ADDobody-AH0 and ADAH11 co-eluting at the expected molecular weight (∼50 kDa) (Fig. S7B,C), evidencing complex formation. A binding constant (K_D_) of ∼82 nM was determined by SPR (Fig. 4B). Next, binding of Chimera AH0 to ADAH11 was analyzed. Chimera AH0 was passed at concentrations ranging from 0.325 nM to 1 nM over immobilized ADAH11, revealing strikingly low dissociation kinetics and an estimated avidity in the picomolar range (∼200 pM. Fig. 4C).

**Figure 4.**
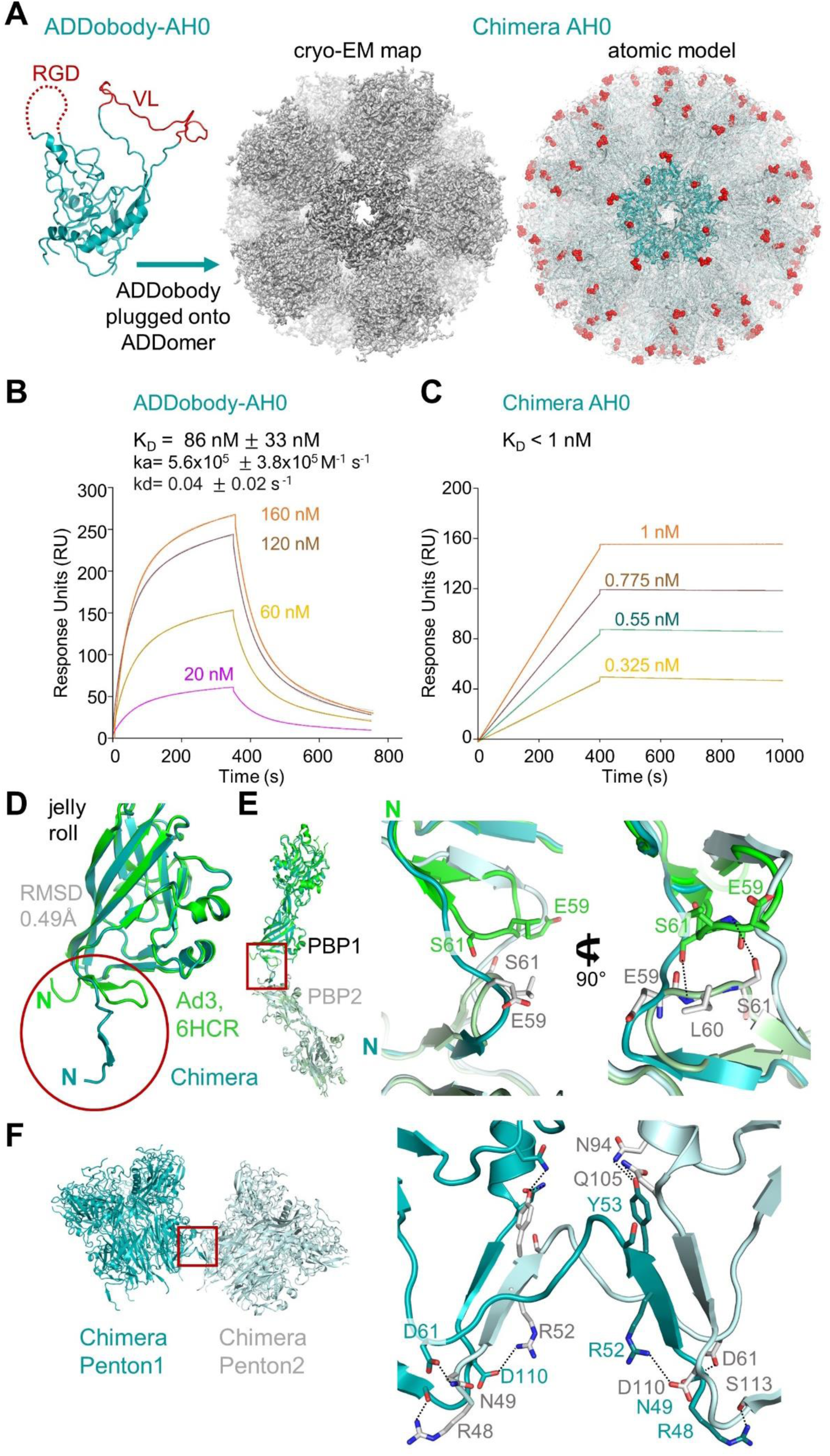
‘ADDomerization’ of ADDobody-AH0. **(A)** Right: ADDobody-AH0 (teal, loops in red) was conjoined with the jelly-roll fold domain of Chimpanzee adenovirus Y25 penton base protein (PBP). Middle: Cryo-EM map of the resulting Chimera AH0. Left: Atomic model of Chimera AH0 (palegreen). One penton is shown in teal, loop residues are shown as red spheres. **(B)** Surface plasmon resonance (SPR) of ADAH11 binding to ADDobody-AH0. Concentrations between 20 nM and 160 nM were flowed over 7,376 RU of ADDobody-AH0 immobilized on a CM5 sensor chip. A 1:1 binding model was used to calculate the *K*D value (82 nM) and standard deviation. **(C)** SPR of Chimera AH0. Concentrations between 0.325 nM and 1 nM were flowed over 1,137 RU of ADAH11 immobilized on a CM5 sensor chip. Fitting indicates a K_D_ in the picomolar range. **(D)** Alignment of the jelly-roll folds of Chimpanzee Y25 adenovirus PBP (teal) and ADDomer Ad3 adenovirus PBP (green, PDB ID: 6HCR ^16^) with an RMSD of 0.371 Å. The N-termini are highlighted (red circle). **(E)** Left: Alignment of PBPs from two adjacent pentons, highlighting the N-terminal region (red box). Middle and right: Close-up views of the N-terminal region, showing loop formation for the Ad3 PBPs (green and palegreen) and strand swapping for the Chimera PBPs (teal and palecyan). In Ad3, the S61 side chains form hydrogen bonds with the E59 peptide backbones. **(F)** Close-up view of the penton-penton interface of Chimera AH0 (red box). An antiparallel β-sheet is formed between residues 50-54 of PBP1 and residues 106-110 of PBP2. Polar contacts formed between side chains are indicated (black dashed lines).

Finally, to validate our Chimera design at near-atomic resolution, we determined the structure by cryo-EM. Highly purified Chimera was used for data collection (Fig. S8 and Table S3). After two-dimensional (2D) and three-dimensional (3D) classification and 3D refinement with applying icosahedral symmetry, we obtained a 2.2 Å resolution structure from 377,978 particles (Fig. 4A, Fig. S8, Fig. S9), resulting in the highest resolution for any ADDomer-derived nanoparticle structure to date. Chimera displays the characteristic overall structure observed previously for the icosahedral ADDomer derived from human adenovirus Ad3 serotype, adopting a dodecamer of pentons (PDB: 6HCR) ^16^. The overlay of the ADDomer Ad3 jelly-roll fold and the Chimpanzee jelly-roll fold domain in our cryo-EM structure reveal a virtually identical fold, with an RMSD of 0.49 Å and differences only found in the N-terminal region of the jelly-roll (Fig. 4D). The N-terminal region of the penton base protein (PBP) is essential for formation of stable dodecameric particles ^18^. In the structure of the Chimera nanoparticle, the N-terminal regions comprising residues G47 to L59 of adjacent PBPs are observed to undergo strand swapping, stabilizing the interaction between pentons (Fig. 4E). Strand swapping has been described before for human adenovirus Ad3 penton base dodecamers ^18^. In contrast, in our previous cryo-EM structure of the Ad3 ADDomer we observed a hairpin conformation of the N-terminal residues, and the interaction between the N-termini was stabilized by hydrogen bonds between S61 and the main chain of E59 (Fig. 4E) ^16^. Notably, N-terminal residues 40-120 of Ad3 ADDomer and 36-116 of Chimera AH0 are identical in these constructs, but the penton-penton interactions are markedly different. In our structure, the penton-penton interface of Chimera AH0, the N-terminus of the PBP forms an anti-parallel β-sheet with a PBP from an adjacent penton, involving residues 50-54 and residues 106-110 (Fig. 4G). In addition, polar contacts are formed between R48 and S113, N49 and D61, R52, while D110 and Y53 can interact with N94 and Q110, further stabilizing the penton-penton interface (Fig. 4G).

In conclusion, we validated our Chimera nanoparticle design and demonstrate that ‘ADDomerization’ of ADDobodies indeed results in picomolar super-binders against a given target.

## Discussion

The ADDomer nanoparticle has been used to develop vaccine candidates for diverse infectious diseases, displaying 60 and more B-cell and T-cell epitopes from SARS-CoV-2 ^17,20^, Chikungunya virus ^16^ or Type O foot-and-mouth disease ^30^. Here, we engineered ADDobody in order to expand the scope of possible applications of the ADDomer nanoparticle, by converting it into a readily customizable, high-avidity super-binder against a target of choice. ADDobody is derived from the crown domain of the penton base protein, the protomer that self-assembles into ADDomer. Our ADDobody design retains the two hyper-variable VL and RGD loops that can be exploited for multi-epitope display for instance to yield vaccines, or, akin to antibody complementarity-determining regions, for recognizing and tightly binding a target molecule. We show that ADDobody can be expressed in *E. coli* at yields comparable to other scaffolds, is monomeric and retains the thermostability observed for ADDomer (Fig. 1C,E). We solved crystal structures of our ADDobody prototype at 2.9 Å and 3.2 Å resolution. Our crystal structures confirm that ADDobody adopts the crown domain fold (Fig. 2A,B) which is unique to adenovirus capsid protein according to the DALI protein structure comparison server ^31^. Strikingly, ADDobodies form pentamers in the crystals indicating ‘molecular memory’, implying that they actually may contribute to pentamerization of the PBPs which was thought to be mediated by the jelly-roll fold. In the crystals, ADDobody penton rings can stack on top of each other, forming decameric barrel shapes (Fig. 2D,F) or form individual pentameric rings interacting with neighboring pentons via coordinating zinc ions (Fig. 2G). Importantly, we show that ADDobodies can be used for ribosome display *in vitro* selections to enrich binders over non-binders (Fig. 3). We estimate an average enrichment of ∼10,000-fold per round of ribosome display for ADDobody-AH0. Similar experiments have been performed with single-chain variable domain (scFv) antibodies previously, to show that ribosome display is a *bona fide in vitro* selection method for binders ^32^. Subsequently, it was shown that picomolar binders can be selected and evolved *in vitro* starting from a synthetic scFv library with a size of ∼10^9^ members ^3^. We confirmed that we can convert ADDobody-AH0 into an ADDomer nanoparticle by fusing the crown domain with a jelly-roll domain from a chimpanzee adenovirus, as an example how ADDobody binders against a target of choice can be selected by ribosome display, and then converted into multimeric ultra-high avidity superbinders. The resulting Chimera nanoparticle assembled correctly and showed indeed high avidity as compared to monomeric ADDobody (Fig. 4B,C), confirming the concept. The Chimera structure surprisingly showed stabilization of the dodecamer by N-terminal strand swapping between pentons, while our previous Ad3 ADDomer was stabilized by hairpin interaction ^16^ (Fig. 4E,F). This finding cannot be explained by the amino acid residues directly involved in penton-penton interaction given that the sequences are identical in both. We speculate that long-range allosteric effects caused by the heterologous crown domain, derived from Ad3, subtle differences at the junctions between the crown domain and the jelly-roll fold, and the different VL and RGD loop sequences contribute to the distinct penton-penton interactions we observed.

We and others have shown that a range of peptides and small protein domains can be inserted into the VL and RGD loops of the PBP ^16,17,19,20,30^. Moving forward, based on our results, we can now conceivably generate a synthetic ADDobody library with randomized VL and RGD loops that can be used for ribosome display *in vitro* selection and evolution experiments against any target of choice. The key advantage of ADDobody as a binder scaffold is that, by rejoining it with a jelly-fold roll domain, it can be readily converted back into a functional PBP which then self-assembles into the icosahedral nanoparticle comprising 12 pentons, for a total of 60 PBPs and thus 60 crown domains per nanoparticle. We demonstrated this here using an unrelated Chimpanzee jelly-roll fold domain to generate Chimera AH0 from our human Ad3 derived ADDobody-AH0 (Fig. 4A). A 2.2 Å resolution cryo-EM structure confirmed successful assembly of the chimeric PBP into a dodecamer of pentons compellingly validating our design (Fig. 4A, Fig. S7, S8, S9). This ‘ADDomerization’ generates 60 binding sites on the nanoparticle resulting in ultra-high avidity even when starting from moderate binding affinities - ADDobody-AH0 binds its ADAH11 target with a K_D_ of ∼82 nM, while Chimera AH0 binds ADAH11 with picomolar avidity with extremely slow dissociation kinetics, impressively illustrating the effect of 60 adjacent binding sites on a single particle.

How could such a super-binder conceivably make a difference in the future? Ultra-high affinity/avidity is required for detection of toxins or chemicals, and for efficient clearance of toxins from organisms. We are particularly interested in snake toxins, and in innovating snakebite treatments, for use in particular in regions that are remote and do not afford a reliable cold chain. These regions are located mostly in low- and middle-income countries (LMICs). Deployment of thermostable ADDomer-based super-binders, that can be transported and stored independent of a cold chain, could provide a useful means over conventional IgG-based antivenoms in tropical areas where snakebite envenoming is a significant health, social and economic burden. The World Health Organization declared snakebite envenoming as a neglected tropical disease and listed the generation of safe, effective and affordable treatment as a key objective ^33,34^. Based on our foundational work presented here, we anticipate that specific anti-toxin ADDobodies could be selected from a synthetic ADDobody library and converted into thermostable ADDomer nanoparticles, to provide a basis of next-generation antivenom treatment.

## STAR Methods

### KEY RESOURCES TABLE

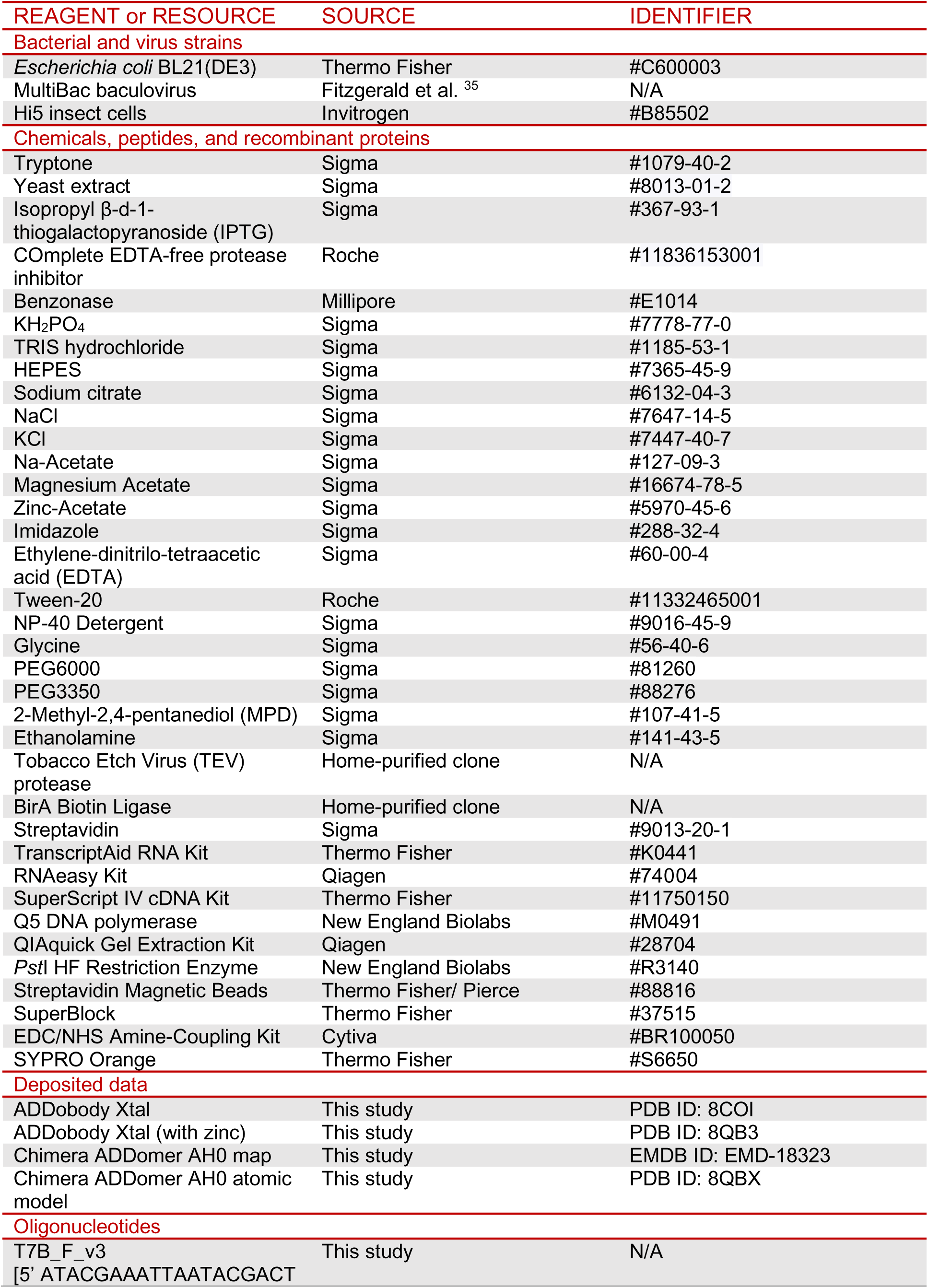
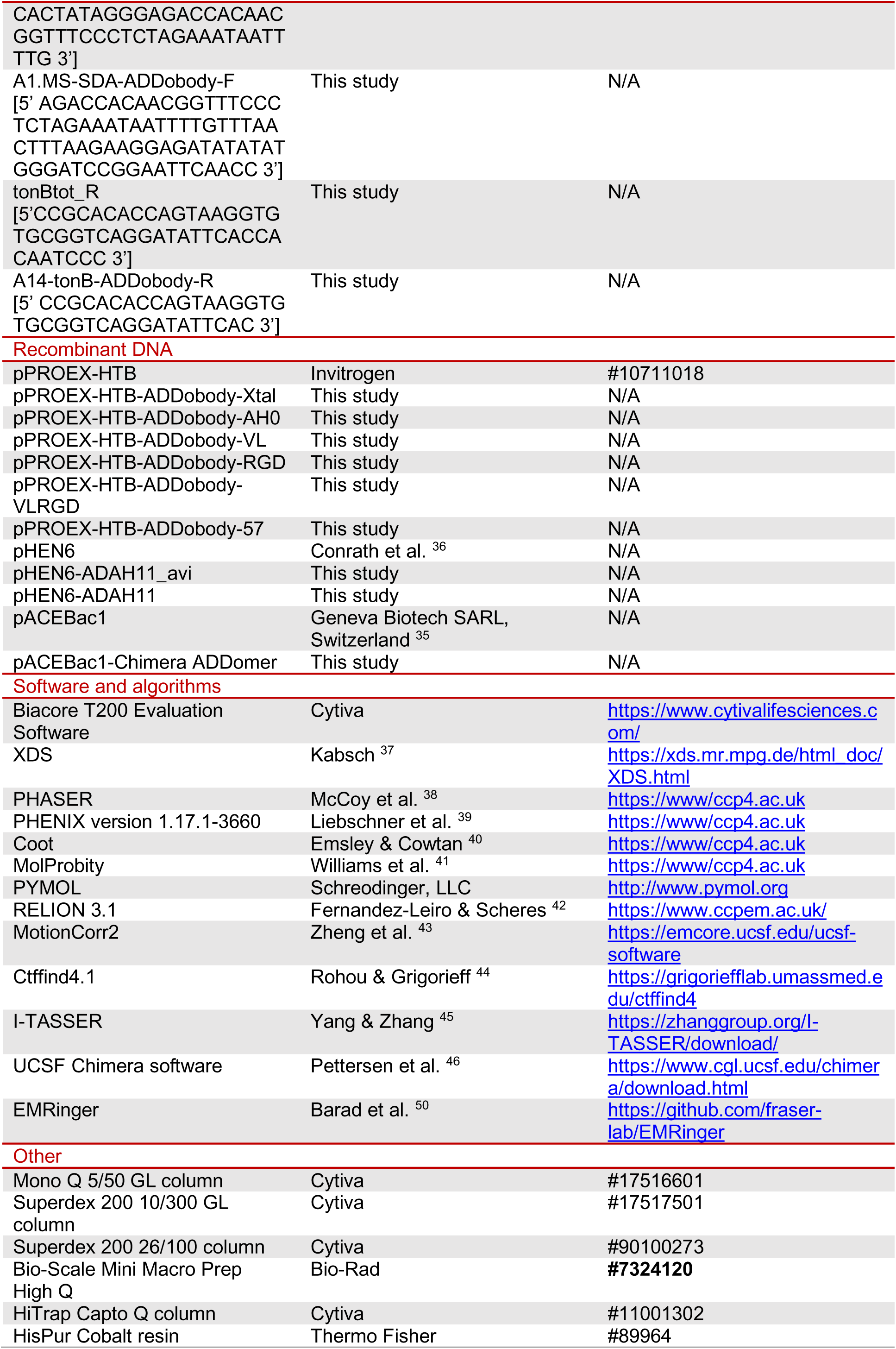
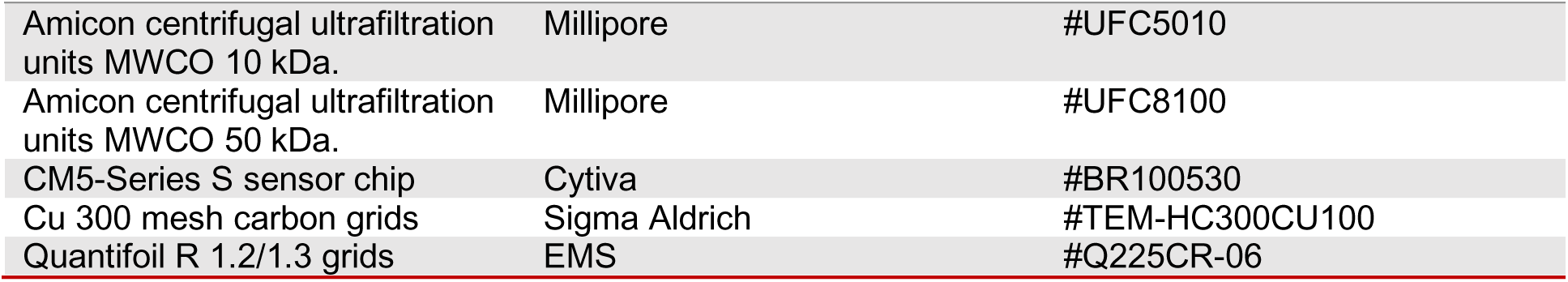

### Lead contact

Further information and requests for resources and reagents should be directed to the lead contact, Christiane Schaffitzel.

### Materials availability

The plasmids generated in this study are available from the lead contact upon request.

### Data and code availability

- All X-ray diffraction data, the EM map and model have been deposited in the PDB and EMDB. They are publicly available as of the date of publication. Accession numbers are listed in the key resources table.
- This paper does not report original code.
- Any additional information required to reanalyze the data reported in this paper is available from the lead contact upon request.

## Method Details

### Construct design and cloning

Based on structural data of the Ad3 adenovirus PBP (PDB: 4AQQ), a DNA sequence encoding the crown domain encompassing residues 132 - 461 was codon-optimized for *E.coli* expression, and a 6 residue long, flexible linker encoding NGDSGN sequence was introduced to reconnect the crown domain replacing two β-strands from the jelly-roll fold domain (Fig. 1B). ADDobody constructs were gene-synthesized (Genscript) and ligated into the pPROEX-HTB vector (Invitrogen) using *Eco*RI and *Not*I restriction sites. ADDobody-Xtal construct was optimized for crystallization by minimizing the VL and RGD flexible loop regions from 12 to 5 residues and 45 to 4 residues, respectively. The ADDobody-AH0 construct contained the SARS-CoV-2 Spike protein RBM-derived AH0 sequence YQAGSTPCNGVEGFNCYFPLQSYGFQPTNGVGY, inserted into the VL loop region ^17^. To facilitate insertion of sequences into the loop regions, we adopted a BioBrick design with restriction sites marking the boundaries of the VL loop (*Eco*RI and *Rsr*II) and the RGD loop (*Bss*HII and *Xba*I) (Fig. 1) ^16^. The amino acid sequences of all ADDobodies used in this study are provided in Supplementary Table S1.

The Chimera construct was designed to comprise the crown domain representing ADDobody and the jelly-roll domain of the chimpanzee derived adenovirus Y25 PBP (UniProt ID: G9G849) comprising an A57S mutation. Furthermore, we implemented mutations in the Chimera nanoparticle to conserve the metal-ion cluster we identified previously in the human Ad3 derived ADDomer ^16^. The resulting gene encoding for the Chimera PBP was synthesized (Genscript), and inserted into plasmid pACEBac1 (Geneva Biotech SARL, Switzerland) using *Bam*HI and *Hind*III restriction sites, giving rise to pACEBac-Chimera.

### ADDobody expression and purification

Plasmids encoding the individual ADDobody constructs were transformed into *E. coli* BL21(DE3) (Thermo Fisher) and grown in 2xYT media. At an OD_600nm_ of 0.9, protein expression was induced with 1 mM isopropyl β-d-1-thiogalactopyranoside (IPTG) for 3 hours at 30 °C with agitation. The cells were harvested by centrifugation at 4000 x g for 15 min at 4 °C. The supernatant was removed, and the pellets were flash-frozen in liquid nitrogen, followed by storage at −80 °C.

For purification, cell pellets from 1 litre culture were resuspended in 20 ml 1x phosphate-buffered saline (PBS), lysed via sonication for 10 minutes with pulse settings (10 s on, 15 s off) at 70% amplitude while lysate was chilled on ice, and the lysate clarified at 40,000 x g for 15 min at 4 °C. The supernatant was applied to 2 ml HisPur Cobalt resin (Thermo Fisher) and incubated for 16 hours at 4 °C while gently mixed by rotation. After washing the resin with high-salt wash buffer (50 mM Tris, 1 M KCl, 5 mM imidazole pH 7.4) and 1x PBS, proteins were eluted using 1x PBS buffer with 200 mM imidazole. Eluted protein-containing fractions were dialyzed into 1x PBS with 2 mM beta-mercapto-ethanol while incubated with 0.5 mg Tobacco Etch Virus (TEV) protease for the removal of the 6xHis-tag. The proteins were further purified by reverse immobilized metal ion affinity chromatography (IMAC) using TALON resin (Qiagen), to remove uncleaved protein, followed by ion exchange purification using a 5 ml Mono Q 5/50 GL column (Cytiva) equilibrated in Buffer A (1x PBS, pH 7.4) and proteins were eluted by using gradient buffer B (from 0.1 M to 1 M NaCl in 1x PBS, pH 7.4). Subsequently, size-exclusion chromatography (SEC) with a Superdex 200 10/300 GL column (Cytiva) equilibrated with 1x PBS, pH 7.4 was used to further purify proteins. Proteins were concentrated using Amicon centrifugal ultrafiltration units with a molecular weight cut-off of 10 kDa. Protein concentrations were determined by measuring UV absorbance at 280 nm with calculated molecular weights and extinction coefficients based on ProtParam EXPASY tool (https://web.expasy.org/protparam/) using a NanoDrop One spectrophotometer (Thermo Fisher). Aliquots of purified protein in 1x PBS, pH 7.4 were flash-frozen in liquid nitrogen and stored at −80 °C.

### Chimera nanoparticle expression and purification

Chimera was expressed using the MultiBac baculovirus expression system following established protocols ^47^ and purified as described previously ^17^. Briefly, pellets were resuspended in Resuspension Buffer (50 mM Tris pH 7.5, 150 mM NaCl, 2 mM MgCl_2_, 1 ml per 2.5×10^7^ cells) supplemented by EDTA-free complete protease inhibitor (Roche). Lysate was prepared by three cycles of freeze-thawing, cleared by centrifugation (40,000g, 30 min), supplemented with Benzonase (Sigma-Aldrich) and incubated on ice for 2 hours. Precipitate was removed by centrifugation (4000 x g, 15 min), the supernatant passed through a 0.45 µm filter, and subjected to size exclusion chromatography (SEC) using a XK 26/100 column (GE Healthcare) equilibrated with 50 mM Tris pH 7.5, 150 mM NaCl running buffer. Fractions containing Chimera were pooled and further purified by ion exchange chromatography (IEX) using a 5 ml Bio-Scale Mini Macro Prep High Q (Bio-Rad) equilibrated in Buffer A (50 mM Tris pH 7.5, 150 mM NaCl) and a linear salt gradient from 0.15 M to 1 M NaCl. Highly purified Chimera eluted at ∼250 mM to 400 mM NaCl. Fractions were pooled, concentrated using Amicon centrifugal ultrafiltration units with a molecular weight cut-off of 50 kDa. Aliquots of purified protein in buffer containing 50 mM Tris pH 7.5, 150 mM NaCl were flash-frozen in liquid nitrogen and stored at −80 °C.

### ADDobody proof-of-concept ribosome display selection

Ribosome display selections were carried out as described previously ^28^. Briefly, the gene fragments encoding ADDobody-AH0 and ADDobody-57 were converted into ribosome display format by adding a C-terminal spacer sequence derived from the periplasmic part of tonB from *E. coli* ^3^ and N-terminal ribosome binding site. 1 µg of ADDobody-AH0 and ADDobody-57 DNA in ribosome display format were transcribed *in vitro* using the TranscriptAid RNA kit (Thermo Fisher). The mRNA was purified using the RNAeasy kit (Qiagen). *E. coli* cell extract for *in vitro* translation was prepared as described ^48^. ADDobody-AH0 RNA was diluted into ADDobody-57 RNA using different ratios (1,1:1, 1:10^3^, 1:10^5^, 1:10^7^, and 1:10^9^). A total of 10 μg RNA mixtures was used in *in vitro* translation reactions which were performed as described ^3^. Next, translation products in solution were incubated with 100 nM biotinylated antigen (ADAH11-Bio). In negative control reactions, no antigen was added in translation products which contain only ADDobody-AH0 or ADDobody-57. After 60 min of incubation at 4 °C, the antigens were captured for 15 mins at 4 °C with 50 µl Streptavidin Magnetic beads (Thermo Fisher) which were previously incubated with superblock (Thermo Fisher) for around 1 h. After 6 washing steps (50 mM Tris-acetate, 150 mM NaCl, 50 mM magnesium acetate, 0.1% Tween-20, pH 7.5 at 4 °C), the mRNA was eluted with EDTA-buffer (50 mM Tris-acetate, 150 mM NaCl, 25 mM EDTA, pH 7.5 at 4°C) and purified using the RNAeasy kit (Qiagen).

Subsequently, the mRNA was immediately reverse transcribed using the SuperScript IV cDNA kit (Thermo) and primer tonBtot_R (Table S4). The resulting cDNA was PCR amplified using primers A1.MS-SDA-ADDobody-F and tonBtot_R (Table S4) using Q5 DNA polymerase (New England Biolabs) and reactions conditions according to the manufacturer’s recommendations (initial denaturation at 98 °C for 30 s, followed by 30 cycles of: 98 °C for 10 s, 68 °C for 20 s, 72 °C for 30 s, and a final extension at 72 °C for 2 mins). Then, the PCR product was gel-extracted using QIAquick Gel Extraction Kit (Qiagen). This was followed by a second PCR amplification step with primers T7B_F_v3 and A14-tonB-ADDobody-R (same PCR program as described above) (Table S4). The final PCR product was gel extracted and used for the next round of *in vitro* transcription and selection.

A unique restriction site (*Pst*I) is present in the gene fragment encoding ADDobody-AH0 (in ribosome display selection format), but not in the ADDobody-57 construct. Therefore, *Pst*I was used to digest the PCR product after ribosome display selection rounds to test if the AH0 construct was enriched. *Pst*I digestion of AH0 generates 2 fragments (242 bp and 1390 bp) (Fig. 3A). 1 µg of purified DNA (PCR product after 2nd round PCR after selections) was incubated with *Pst*I-HF (New England Biolabs) at 37 °C for 1 h. Subsequently, the reaction was analyzed by agarose gel electrophoresis.

### Expression, purification, and biotinylation of ADAH11

The gene encoding Avi-tagged ADAH11 (ADAH11-Avi) was cloned into the plasmid pHEN6 ^36^, which encodes a C-terminal 3xFLAG and 6xHis tag and an N-terminal pelB signal sequence for the expression of proteins in the bacterial periplasm. For expression, the plasmid pHEN6_ADAH11 was transformed into the *E. coli* BL21(DE3) strain. Cells were grown in 2xYT media at 37 °C until OD_600_ around 0.8, and protein expression was induced by 1 mM IPTG at 30 °C overnight. Cells were harvested by centrifugation (3200xg, 15 min) and, to extract the bacterial periplasm, cell pellets were resuspended in 10 ml cold TES buffer (50 mM Tris pH 8.0, 20% Sucrose, 1 mM EDTA) supplemented with EDTA-free complete protease inhibitor (Roche, Switzerland) and incubated at 4 °C for 45 mins. Subsequently, 15 ml of ice-cold shock buffer (20 mM Tris pH 8.0, 5 mM MgCl_2_) was added and incubated at 4 °C for 45 mins, followed by centrifuging at 13,000xg for 30 mins at 4 °C. The supernatant was kept as the periplasm. The supernatant containing the periplasm was applied to 2 ml HisPur Cobalt resin (Thermo Fisher) and incubated for 2 hours at 4 °C. After washing the resin with wash buffer 1 (50 mM HEPES, 200 mM KCl, 10 mM imidazole pH 8.0) and wash buffer 2 (50 mM HEPES, 200 mM KCl, 20 mM imidazole pH 8.0), the fractions of ADAH11-Avi were eluted in elution buffer (50 mM HEPES, 200 mM KCl, 500 mM imidazole pH 8.0). The purified protein was dialyzed into PBS, pH 7.4, at 4 °C overnight (Fig. S5).

In order to biotinylate ADAH11-Avi, 30 μM ADAH11-Avi protein was incubated with 2 μM maltose binding protein-tagged biotin ligase BirA in PBS buffer containing 5 mM MgCl_2_, 5 mM ATP, and 150 μM biotin at 4 °C overnight. Afterwards, size exclusion chromatography was performed using a Superdex 75 pg column (GE Healthcare Life Sciences) to remove excess BirA protein, biotin and to further purify ADAH11-Avi. Anion exchange chromatography (HiTrap Capto Q column) was conducted to further purify biotinylated ADAH11 and to remove any RNases using PBS pH7.4 buffer and applying a linear salt (from 0.1 M to 1 M NaCl) gradient. Biotinylated ADAH11-Avi (ADAH11-Bio) eluted at ∼400 mM NaCl.

To determine the biotinylation efficiency, a gel shift assay using streptavidin (Sigma) was performed. Biotinylated ADAH11 was pre-incubated with streptavidin and loaded onto an SDS-gel, resulting in a shift of the ADAH11-Bio protein to higher molecular weight. As a control, a sample without streptavidin was loaded onto the same SDS-gel (Fig. S5).

### Surface plasmon resonance assays

Surface plasmon resonance (SPR) experiments were performed on a Biacore T200 using a CM5-Series S sensor chip (Cytiva). Freshly prepared, filtered, and degassed HBS-P+ running buffer (10 mM HEPES pH 7.4, 150 mM NaCl and 0.05% v/v NP40) was used at 25 °C. ADDobody-AH0 was immobilized by 1-Ethyl-3-[3-dimethylaminopropyl] carbodiimide hydrochloride (EDC) / N-hydroxy succinimide (NHS) chemistry, using an amine-coupling kit (Cytiva). A flow cell pair (Fc1 and Fc2) was activated using the amine coupling kit, and immobilization of the protein was carried out on a single flow cell leaving a second flow cell as a background control for signal subtraction. The chip surface was activated by injecting a 1:1 (v/v) mixture of 200 mM EDC and 50 mM NHS for 7 min. 30 μg purified ADDobody-AH0 in 10 mM Na-acetate pH 4.5 at a concentration of 300 μg/ml was injected for 10 min, followed by injection of 100 mM ethanolamine pH 8.5 for 1 min leading to surface inactivation. After immobilization, ∼7400 response units (RU) were reached.

ADAH11 binding to ADDobody-AH0 was measured using concentrations ranging from 20 nM to 160 nM in HBS-P+ running buffer. Samples were injected at 30 µl/min with an association phase of 400 s and a dissociation phase of 400 s. After every experiment, the surface was regenerated with a 60 s injection regeneration solution (10 mM glycine pH 2.0). For Chimera AH0 binding experiments, 25 µg purified ADAH11 in 10 mM Na-acetate pH 4.5 at a concentration of 250 μg/ml was immobilized as outlined above for ADDobody-AH0. After immobilization, ∼1100 response units (RU) were reached. Chimera AH0 binding to ADAH11 was tested using concentrations ranging from 0.325 nM to 1 nM in HBS-P+ running buffer. Samples were injected at 30 µl/min with an association phase of 400 s and a dissociation phase of 600 s. After every experiment, the surface was regenerated with a 60 s injection of regeneration solution (10 mM glycine pH 1.5). Sensorgrams were analyzed with the Biacore Evaluation software (version 1.0) yielding on- and off-rates obtained through fitting the association and dissociation phases of four different concentrations of each analyte, each measured 3 times (n=3), using the one-to-one binding model.

### Thermal shift assays

Thermal shift experiments on highly purified ADDobody constructs were performed using a ThermoFluor assay ^27^. ADDobody samples were diluted to 1 mg/ml in 1 x PBS buffer and supplemented with 1 x SYPRO Orange probe in 25 µl reaction volumes (Thermo Fisher). Samples were subjected to thermal denaturation in a Real Time PCR machine (Agilent Stratagene Mx3005P). Fluorescence intensity of the probe was recorded throughout denaturation using a temperature gradient from 25 °C to 95 °C, with a step size of 1 °C per minute, using excitation and emission wavelengths of 492 nm and 516 nm, respectively. Relative fluorescence emission intensity (R) was plotted as a function of the temperature and normalized to range between 0 to 1. The melting temperature (Tm) was determined as the temperature corresponding to the midpoint between the baseline and the point with maximum fluorescence intensity.

### Protein crystallization

Crystallization experiments were performed with ADDobody-Xtal using a standard sitting drop vapor diffusion method on MRC 2 Well Crystallization Polystyrene plate (Swissci) containing 60 µl of reservoir solution. Using a Mosquito crystallization robot (SPT Labtech) 150 nl and 175 nl of 5 mg/ml ADDobody-Xtal in 1x PBS buffer was dispensed along with 150 nl and 125 nl of reservoir solution in drops A and B respectively resulting in 300 nl total drop volume. Plates were incubated at 20 °C. The best diffracting crystals appeared after 7 days in 20 % (w/v) PEG6000, 0.1 M zinc-acetate, 0.1 HEPES pH 7.0 for zinc-containing ADDobody; and after 10 days in 20% (w/v) PEG3350, 0.15 M citrate pH 5.5 for zinc-free ADDobody. Crystals were transferred in a solution containing 20% 2-Methyl-2,4-pentanediol (MPD) as a cryoprotectant in the crystallization condition prior to flash freezing in liquid N_2_.

### Data collection and structure determination

Diffraction data were collected on the I03 (zinc-free ADDobody) and I04 beamlines (zinc-containing ADDobody) at the Diamond Light Source under nitrogen cryo-stream (∼100 K) (Harwell Science and Innovation Campus) and images processed using XDS (version 03/2019) and scaled with AIMLESS ^37^ (Table S2). Structures were solved by molecular replacement in PHASER using the Ad3 PBP crown domain structure (PDB ID: 4AQQ) as an input model ^16^. Manual model building was performed in Coot (version 0.8.9.2) and refinement carried out in PHENIX.REFINE (PHENIX version 1.17.1-3660) ^39,40^. Models were validated and statistics obtained using MolProbity ^41^. Figures were prepared using PyMol (version 4.6.0).

### Negative-stain EM

Negative-stain EM quality control was performed with purified Chimera (0.1 mg/ml) dialyzed into 50 mM TRIS (pH 7.5), 150 mM NaCl. 4 µl sample was applied onto glow discharged Cu 300 mesh carbon grids before staining with 2% (w/v) uranyl acetate. Images were acquired with a 120 kV FEI Tecnai 12 electron microscope equipped with a Ceta camera (Thermo Fisher).

### Cryo-EM and data collection

4 μl of purified Chimera (0.5 mg/ml) was applied onto carbon-coated Quantifoil R 1.2/1.3 grids (Sigma-Aldrich) which previously were glow discharged for 120 seconds at 5 mA. Excess protein was blotted away for 2 seconds at 4 °C in 100% relative humidity before being plunge frozen into liquid ethane using a Vitrobot Mark IV (Thermo Fisher). Cryo-EM data were collected with a 200 kV FEI Talos Arctica microscope equipped with a Gatan K2 direct electron detector and an energy filter using automated acquisition software (EPU). In total, 6,179 dose-fractioned movies were recorded in super-resolution mode each containing 40 frames with an accumulated total dose of 42.4 e/Å^2^ recorded in super-resolution mode at a nominal magnification of 130,000x corresponding to a physical pixel size of 1.05 Å and a virtual pixel size of 0.525 Å. Images were recorded with a defocus range of −0.8 to −2.0 μm (Table S3).

### Image processing

Image processing was performed with the RELION 3.1 software package ^42^. The micrographs were motion-corrected using MotionCorr2 ^43^, and contrast transfer function (CTF) information was determined using ctffind4.1 ^44^. Micrographs with significant astigmatism (> 500) and defocus (> 2.1 μm) were removed. This resulted in the selection of 5,474 micrographs with CTF rings extending below 4.0 Å that were used for further processing. A total of 710,760 particles were boxed using RELION autopicking software. Several rounds of reference-free 2D classification were performed (Fig. S8) yielding 566,795 particles which were subjected to 3D classification. Class 1 (of two) comprised 377,978 good particles which were subjected to 3D refinement yielding a reconstruction of ∼2.6 Å resolution. Particles were further subjected to Bayesian polishing and CTF refinement and final 3D refinement with imposed icosahedral symmetry. Subsequently, postprocessing was performed for masking and automatic B-factor sharpening with a B-factor value of −88. The resolution of the final map was determined to be 2.2 Å based on the Fourier shell correlation (FSC) 0.143 cutoff criterion ^49^ (Fig. S9A). Local resolution was calculated using the local resolution estimation program in RELION (Fig. S9B). Image processing and 3D classification of Chimera nanoparticle were performed using public cloud resources provided by the Oracle Cloud Infrastructure.

### Model building and refinement

Model building was performed using I-TASSER ^45^ with the Ad3-derived ADDomer structure (PDB ID: 6HCR) ^16^ as template. The generated model was manually docked into the density map in UCSF Chimera software ^46^. Refinement was performed using Phenix Real-Space refinement software version 1.17 ^39^ and COOT ^40^ for model-building, before evaluating the model using MolProbity ^41^ and EMRinger ^50^. Refinement statistics are summarized in Table S2 and S3.

## SUPPLEMENTAL INFORMATION

Supplemental information contains figures S1-S9 and tables S1-S4.

## Supporting information

Supplemental information contains figures S1-S9 and tables S1-S4.

## ACKNOWLEDGEMENTS

We thank all members, present and past, of the Berger and Schaffitzel laboratories for their contributions and helpful discussions. We are grateful for support from the Oracle Higher Education and Research program to enable cryo-EM data processing using Oracle’s high-performance public cloud infrastructure (https://cloud.oracle.com/en_US/cloud-infrastructure), and we thank Simon Burbidge, Christopher Woods, Matt Williams and Richard Pitts for computation infrastructure support. This work was carried out using the computational and data storage facilities of the Advanced Computing Research Centre, University of Bristol. The authors thank University of Bristol and the Max Planck Gesellschaft (MPG), Germany, for generous support through the Max Planck Bristol Centre for Minimal Biology (MPBC). We acknowledge the staff of the BrisSynBio BioSuite including Peter Wilson and the beamline scientists at the Diamond Light Source (via BAG access) for their assistance with crystallographic studies. For the purpose of Open Access, the authors have applied a CC BY public copyright license to any Author Accepted Manuscript version arising from this submission.

We acknowledge support and assistance by the Wolfson Bioimaging Facility and the GW4 Facility for High-Resolution Electron Cryo-Microscopy funded by the Wellcome Trust (202904/Z/16/Z and 206181/Z/17/Z) and BBSRC (BB/R000484/1). I.B. acknowledges support from the EPSRC Future Vaccine Manufacturing and Research Hub (EP/R013764/1) and the ERC (AdvG 834631). C.S. and I.B. are Investigators of the Wellcome Trust (210701/Z/18/Z; 106115/Z/14/Z, 221708/Z/20/Z). C.S. acknowledges funding by the Wellcome Trust – University of Bristol institutional Translation Partnership (iTP, fEC 282839). C.S. and I.B. are supported by a Horizon 2020 FET OPEN grant ‘ADDovenom’ (contract nr. 899670).

## AUTHOR CONTRIBUTIONS

I.B. and C.S. conceived the study. D.B, H.S, I.B. and C.S. prepared figures and wrote the manuscript with help of all coauthors. D.B purified ADDobody-Xtal and Chimera, performed crystallization trials and solved the ADDobody-Xtal structure. D.B. processed cryo-EM data, solved the Chimera, and performed Biacore measurements with help from C.T. H.S. performed ribosome display selections, purifications and thermofluor assays of ADDobodies. R.B.S. computationally designed the ADDobody prototype. F.G. and I.B. designed the Chimera construct. F.G. purified ADDobody and established the Chimera production protocol. J.C. purified Chimera. K.G. supported protein purification, crystallization and data collection from zinc free ADDobody crystals. J.B. and S.Y. supported image processing of Chimera. C.T. supported crystallization, data collection, structure solution, model building and refinement of the X-ray and cryo-EM model. G.G. supported Biacore measurements. U.B. performed cryo-EM data collection.

## DATA AVAILABILITY

All data to evaluate the conclusions in the paper are present in the paper and/or the Supplementary Materials. All datasets generated during the current study have been deposited in the Electron Microscopy Data Bank (EMDB) under accession number EMD-18323 (Chimera) and the Protein Data Bank (PDB) under accession numbers PDB ID 8QBX (Chimera), 8COI (zinc-free ADDobody) and 8QB3 (zinc-containing ADDobody). Reagents are available from C.S. and I.B upon request.

## INCLUSION AND DIVERSITY

We support inclusive, diverse, and equitable conduct of research.

## REFERENCES

1. Winter, G., Griffiths, A.D., Hawkins, R.E., and Hoogenboom, H.R. (1994). Making antibodies by phage display technology. Annu. Rev. Immunol. 12, 433–455. 10.1146/annurev.iy.12.040194.002245.

2. Gai, S.A., and Wittrup, K.D. (2007). Yeast surface display for protein engineering and characterization. Curr. Opin. Struct. Biol. 17, 467–473. 10.1016/j.sbi.2007.08.012.

3. Hanes, J., Schaffitzel, C., Knappik, A., and Plückthun, A. (2000). Picomolar affinity antibodies from a fully synthetic naive library selected and evolved by ribosome display. Nat. Biotechnol. 18, 1287–1292. 10.1038/82407.

4. Zimmermann, I., Egloff, P., Hutter, C.A.J., Kuhn, B.T., Brauer, P., Newstead, S., Dawson, R.J.P., Geertsma, E.R., and Seeger, M.A. (2020). Generation of synthetic nanobodies against delicate proteins. Nat. Protoc. 15, 1707–1741. 10.1038/s41596-020-0304-x.

5. Dreier, B., and Plückthun, A. (2018). Rapid Selection of High-Affinity Antibody scFv Fragments Using Ribosome Display. Methods Mol. Biol. 1827, 235–268. 10.1007/978-1-4939-8648-4_13.

6. Li, R., Kang, G., Hu, M., and Huang, H. (2019). Ribosome Display: A Potent Display Technology used for Selecting and Evolving Specific Binders with Desired Properties. Mol. Biotechnol. 61, 60–71. 10.1007/s12033-018-0133-0.

7. Zahnd, C., Spinelli, S., Luginbühl, B., Amstutz, P., Cambillau, C., and Plückthun, A. (2004). Directed *in vitro* evolution and crystallographic analysis of a peptide-binding single chain antibody fragment (scFv) with low picomolar affinity. J. Biol. Chem. 279, 18870–18877. 10.1074/jbc.M309169200.

8. Boder, E.T., Midelfort, K.S., and Wittrup, K.D. (2000). Directed evolution of antibody fragments with monovalent femtomolar antigen-binding affinity. Proc. Natl. Acad. Sci. USA 97, 10701–10705. 10.1073/pnas.170297297.

9. Koenning, D., and Schaefer, J.V. (2021). Alternative Binding Scaffolds: Multipurpose Binders for Applications in Basic Research and Therapy. In: Rüker, F., Wozniak-Knopp, G. (eds) Introduction to Antibody Engineering. Learning Materials in Biosciences. Springer, Cham. 10.1007/978-3-030-54630-4_9.

10. Gebauer, M., and Skerra, A. (2020). Engineered Protein Scaffolds as Next-Generation Therapeutics. Annu. Rev. Pharmacol. Toxicol. 60, 391–415. 10.1146/annurev-pharmtox-010818-021118.

11. Simeon, R., and Chen, Z. (2018). *In vitro*-engineered non-antibody protein therapeutics. Protein Cell 9, 3–14. 10.1007/s13238-017-0386-6.

12. Lofblom, J., Feldwisch, J., Tolmachev, V., Carlsson, J., Stahl, S., and Frejd, F.Y. (2010). Affibody molecules: engineered proteins for therapeutic, diagnostic and biotechnological applications. FEBS Lett. 584, 2670–2680. 10.1016/j.febslet.2010.04.014.

13. Bloom, L., and Calabro, V. (2009). FN3: a new protein scaffold reaches the clinic. Drug Discov Today 14, 949–955. 10.1016/j.drudis.2009.06.007.

14. Plückthun, A. (2015). Designed ankyrin repeat proteins (DARPins): binding proteins for research, diagnostics, and therapy. Annu. Rev. Pharmacol. Toxicol. 55, 489–511. 10.1146/annurev-pharmtox-010611-134654.

15. Goux, M., Becker, G., Gorre, H., Dammicco, S., Desselle, A., Egrise, D., Leroi, N., Lallemand, F., Bahri, M.A., Doumont, G., et al. (2017). Nanofitin as a New Molecular-Imaging Agent for the Diagnosis of Epidermal Growth Factor Receptor Over-Expressing Tumors. Bioconjug. Chem. 28, 2361–2371. 10.1021/acs.bioconjchem.7b00374.

16. Vragniau, C., Bufton, J.C., Garzoni, F., Stermann, E., Rabi, F., Terrat, C., Guidetti, M., Josserand, V., Williams, M., Woods, C.J., et al. (2019). Synthetic self-assembling ADDomer platform for highly efficient vaccination by genetically encoded multiepitope display. Sci. Adv. 5, eaaw2853. 10.1126/sciadv.aaw2853.

17. Buzas, D., Bunzel, H.A., Staufer, O., Milodowski, E.J., Edmunds, G.L., Bufton, J.C., Vidana Mateo, B.V., Yadav, S.K.N., Gupta, K., Fletcher, C., et al. (2023). Antibodies generated *in vitro* and *in vivo* elucidate design of a thermostable ADDomer COVID-19 nasal nanoparticle vaccine. 2023.2003.2017.533092. 10.1101/2023.03.17.533092 BioRxiv.

18. Szolajska, E., Burmeister, W.P., Zochowska, M., Nerlo, B., Andreev, I., Schoehn, G., Andrieu, J.P., Fender, P., Naskalska, A., Zubieta, C., et al. (2012). The structural basis for the integrity of adenovirus Ad3 dodecahedron. PLoS One 7, e46075. 10.1371/journal.pone.0046075.

19. Villegas-Mendez, A., Garin, M.I., Pineda-Molina, E., Veratti, E., Bueren, J.A., Fender, P., and Lenormand, J.L. (2010). *In vivo* delivery of antigens by adenovirus dodecahedron induces cellular and humoral immune responses to elicit antitumor immunity. Mol. Ther. 18, 1046–1053. 10.1038/mt.2010.16.

20. Chevillard, C., Amen, A., Besson, S., Hannani, D., Bally, I., Dettling, V., Gout, E., Moreau, C.J., Buisson, M., Gallet, S., et al. (2022). Elicitation of potent SARS-CoV-2 neutralizing antibody responses through immunization with a versatile adenovirus-inspired multimerization platform. Mol. Ther. 30, 1913–1925. 10.1016/j.ymthe.2022.02.011.

21. Guo, M., Li, J., Teng, Z., Ren, M., Dong, H., Zhang, Y., Ru, J., Du, P., Sun, S., and Guo, H. (2021). Four Simple Biomimetic Mineralization Methods to Improve the Thermostability and Immunogenicity of Virus-like Particles as a Vaccine against Foot-and-Mouth Disease. Vaccines (Basel) 9 10.3390/vaccines9080891.

22. Zochowska, M., Paca, A., Schoehn, G., Andrieu, J.P., Chroboczek, J., Dublet, B., and Szolajska, E. (2009). Adenovirus dodecahedron, as a drug delivery vector. PLoS One 4, e5569. 10.1371/journal.pone.0005569.

23. Ashok, A., Brison, M., and LeTallec, Y. (2017). Improving cold chain systems: Challenges and solutions. Vaccine 35, 2217–2223. 10.1016/j.vaccine.2016.08.045.

24. Besson, S., Vragniau, C., Vassal-Stermann, E., Dagher, M.C., and Fender, P. (2020). The Adenovirus Dodecahedron: Beyond the Platonic Story. Viruses 12. 10.3390/v12070718.

25. Shetty, R.P., Endy, D., and Knight, T.F., Jr. (2008). Engineering BioBrick vectors from BioBrick parts. J. Biol. Eng. 2, 5 10.1186/1754-1611-2-5.

26. Hoffmann, M., Kleine-Weber, H., Schroeder, S., Kruger, N., Herrler, T., Erichsen, S., Schiergens, T.S., Herrler, G., Wu, N.H., Nitsche, A., et al. (2020). SARS-CoV-2 Cell Entry Depends on ACE2 and TMPRSS2 and Is Blocked by a Clinically Proven Protease Inhibitor. Cell 181, 271–280 e278. 10.1016/j.cell.2020.02.052.

27. Dupeux, F., Rower, M., Seroul, G., Blot, D., and Marquez, J.A. (2011). A thermal stability assay can help to estimate the crystallization likelihood of biological samples. Acta Crystallogr. D Biol. Crystallogr. 67, 915–919. 10.1107/S0907444911036225.

28. Schaffitzel, C., Zahnd, C., Amstutz, P., Luginbühl, B., and Plückthun, A. (2005). In Vitro Selection and Evolution of Protein-Ligand Interactions by Ribosome Display. in Protein-Protein Interactions A Molecular Cloning Manual (eds. Golemis, E. & Adams, P.) 517–548 (Cold Spring Harbor Laboratory Press, Cold Spring Harbor, New York, 2005).

29. Mattheakis, L.C., Bhatt, R.R., and Dower, W.J. (1994). An *in vitro* polysome display system for identifying ligands from very large peptide libraries. Proc. Natl. Acad. Sci. USA 91, 9022–9026. 10.1073/pnas.91.19.9022.

30. Luo, C., Yan, Q., Huang, J., Liu, J., Li, Y., Wu, K., Li, B., Zhao, M., Fan, S., Ding, H., and Chen, J. (2022). Using Self-Assembling ADDomer Platform to Display B and T Epitopes of Type O Foot-and-Mouth Disease Virus. Viruses 14. 10.3390/v14081810.

31. Holm, L. (2022). Dali server: structural unification of protein families. Nucleic Acids Res. 50, W210–W215. 10.1093/nar/gkac387.

32. Hanes, J., and Plückthun, A. (1997). *In vitro* selection and evolution of functional proteins by using ribosome display. Proc. Natl. Acad. Sci. USA 94, 4937–4942. 10.1073/pnas.94.10.4937.

33. Harrison, R.A., and Gutierrez, J.M. (2016). Priority Actions and Progress to Substantially and Sustainably Reduce the Mortality, Morbidity and Socioeconomic Burden of Tropical Snakebite. Toxins (Basel) 8. 10.3390/toxins8120351.

34. World Health Organisation, W.H.O. (2021). Snakebite Envenoming. https://www.who.int/news-room/fact-sheets/detail/snakebite-envenoming.

35. Fitzgerald, D.J., Berger, P., Schaffitzel, C., Yamada, K., Richmond, T.J., and Berger, I. (2006). Protein complex expression by using multigene baculoviral vectors. Nat. Methods 3, 1021–1032. 10.1038/nmeth983.

36. Conrath, K.E., Lauwereys, M., Galleni, M., Matagne, A., Frere, J.M., Kinne, J., Wyns, L., and Muyldermans, S. (2001). Beta-lactamase inhibitors derived from single-domain antibody fragments elicited in the *camelidae*. Antimicrob. Agents Chemother. 45, 2807–2812. 10.1128/AAC.45.10.2807-2812.2001.

37. Kabsch, W. (2010). Xds. Acta Crystallogr. D Biol. Crystallogr. 66, 125–132. 10.1107/S0907444909047337.

38. McCoy, A.J., Grosse-Kunstleve, R.W., Adams, P.D., Winn, M.D., Storoni, L.C., and Read, R.J. (2007). Phaser crystallographic software. J. Appl. Crystallogr. 40, 658–674. 10.1107/S0021889807021206.

39. Liebschner, D., Afonine, P.V., Baker, M.L., Bunkoczi, G., Chen, V.B., Croll, T.I., Hintze, B., Hung, L.W., Jain, S., McCoy, A.J., et al. (2019). Macromolecular structure determination using X-rays, neutrons and electrons: recent developments in Phenix. Acta Crystallogr. D Struct. Biol. 75, 861–877. 10.1107/S2059798319011471.

40. Emsley, P., and Cowtan, K. (2004). Coot: model-building tools for molecular graphics. Acta Crystallogr. D Biol. Crystallogr. 60, 2126–2132. 10.1107/S0907444904019158.

41. Williams, C.J., Headd, J.J., Moriarty, N.W., Prisant, M.G., Videau, L.L., Deis, L.N., Verma, V., Keedy, D.A., Hintze, B.J., Chen, V.B., et al. (2018). MolProbity: More and better reference data for improved all-atom structure validation. Protein Sci. 27, 293–315. 10.1002/pro.3330.

42. Fernandez-Leiro, R., and Scheres, S.H.W. (2017). A pipeline approach to single-particle processing in RELION. Acta Crystallogr. D Struct. Biol. 73, 496–502. 10.1107/S2059798316019276.

43. Zheng, S.Q., Palovcak, E., Armache, J.P., Verba, K.A., Cheng, Y., and Agard, D.A. (2017). MotionCor2: anisotropic correction of beam-induced motion for improved cryo-electron microscopy. Nat. Methods 14, 331–332. 10.1038/nmeth.4193.

44. Rohou, A., and Grigorieff, N. (2015). CTFFIND4: Fast and accurate defocus estimation from electron micrographs. J. Struct. Biol. 192, 216–221. 10.1016/j.jsb.2015.08.008.

45. Yang, J., and Zhang, Y. (2015). I-TASSER server: new development for protein structure and function predictions. Nucleic Acids Res. 43, W174–181. 10.1093/nar/gkv342.

46. Pettersen, E.F., Goddard, T.D., Huang, C.C., Couch, G.S., Greenblatt, D.M., Meng, E.C., and Ferrin, T.E. (2004). UCSF Chimera--a visualization system for exploratory research and analysis. J. Comput. Chem. 25, 1605–1612. 10.1002/jcc.20084.

47. Sari-Ak, D., Bufton, J., Gupta, K., Garzoni, F., Fitzgerald, D., Schaffitzel, C., and Berger, I. (2021). VLP-factory and ADDomer((c)): Self-assembling Virus-Like Particle (VLP) Technologies for Multiple Protein and Peptide Epitope Display. Curr. Protoc. 1, e55. 10.1002/cpz1.55.

48. Estrozi, L.F., Boehringer, D., Shan, S.O., Ban, N., and Schaffitzel, C. (2011). Cryo-EM structure of the E. coli translating ribosome in complex with SRP and its receptor. Nat. Struct. Mol. Biol. 18, 88–90. 10.1038/nsmb.1952.

49. Rosenthal, P.B., and Henderson, R. (2003). Optimal determination of particle orientation, absolute hand, and contrast loss in single-particle electron cryomicroscopy. J. Mol. Biol. 333, 721–745. 10.1016/j.jmb.2003.07.013.

50. Barad, B.A., Echols, N., Wang, R.Y., Cheng, Y., DiMaio, F., Adams, P.D., and Fraser, J.S. (2015). EMRinger: side chain-directed model and map validation for 3D cryo-electron microscopy. Nat. Methods 12, 943–946. 10.1038/nmeth.3541.

