## Supplemental information contains figures S1-S9 and tables S1-S4. for "Engineering the ADDobody protein scaffold for generation of high-avidity ADDomer super-binders"

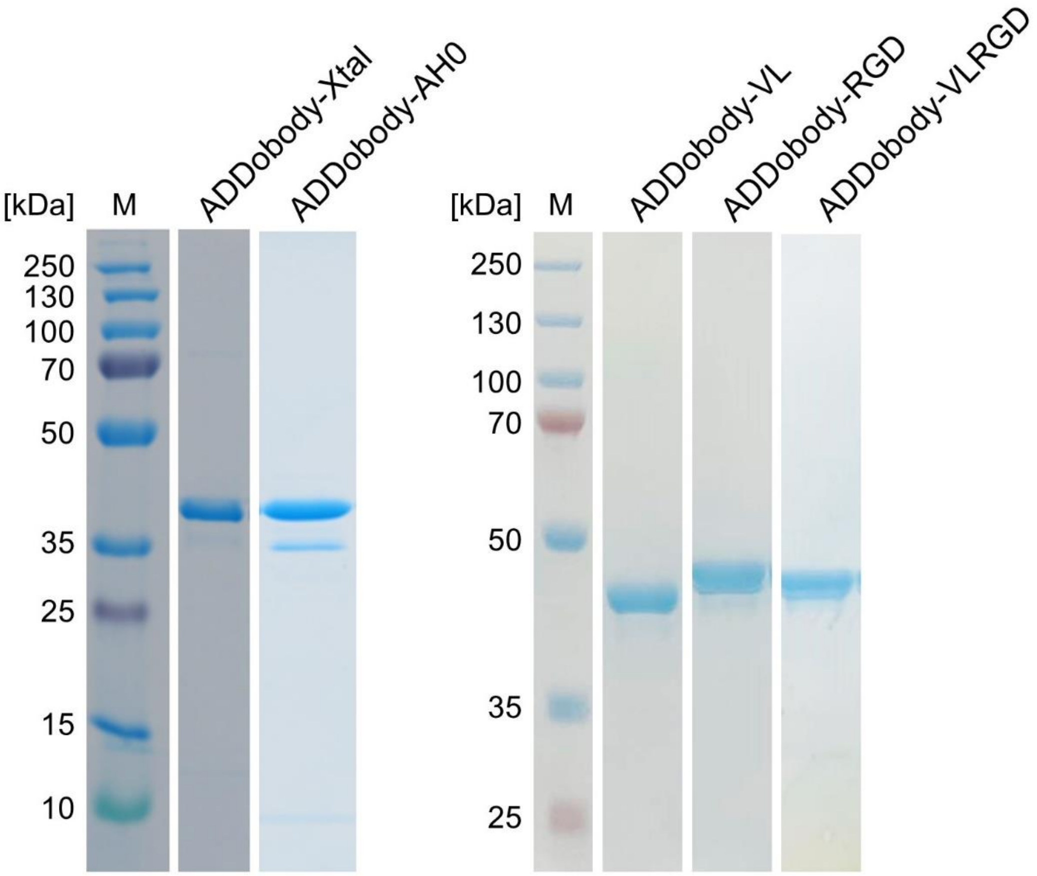


**Figure S1. Coomassie-stained SDS gel showing purified ADDobodies.** Coomassie-stained SDS–PAGE showing purified ADDobody constructs. M: Molecular weight marker. The sequence of the ADDobodies is provided in Supplementary Table S1.

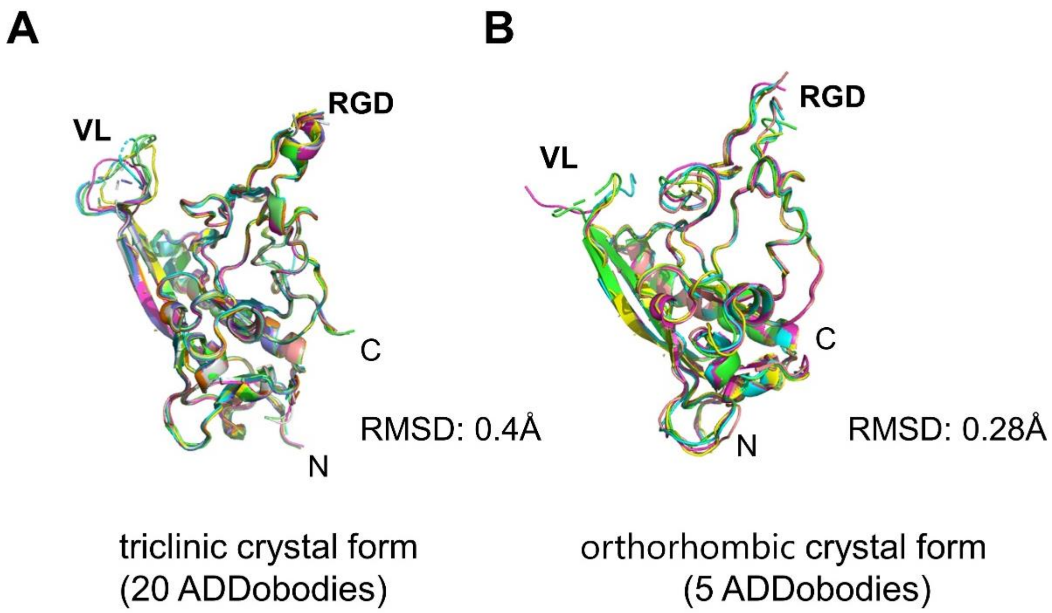


**Figure S2. Superimposition of ADDobodies (A)** from the triclinic (P1) crystal form comprising 20 ADDobody molecules in the asymmetric unit, adopting four pentons forming two decamer barrels, and **(B)** from the $Zn^{2+}$-containing orthorhombic ($P2_12_12_1$) crystal form (5 ADDobodies per asymmetric unit adopting a penton). The root-mean-square deviation (RMSD) of the overlay is indicated.

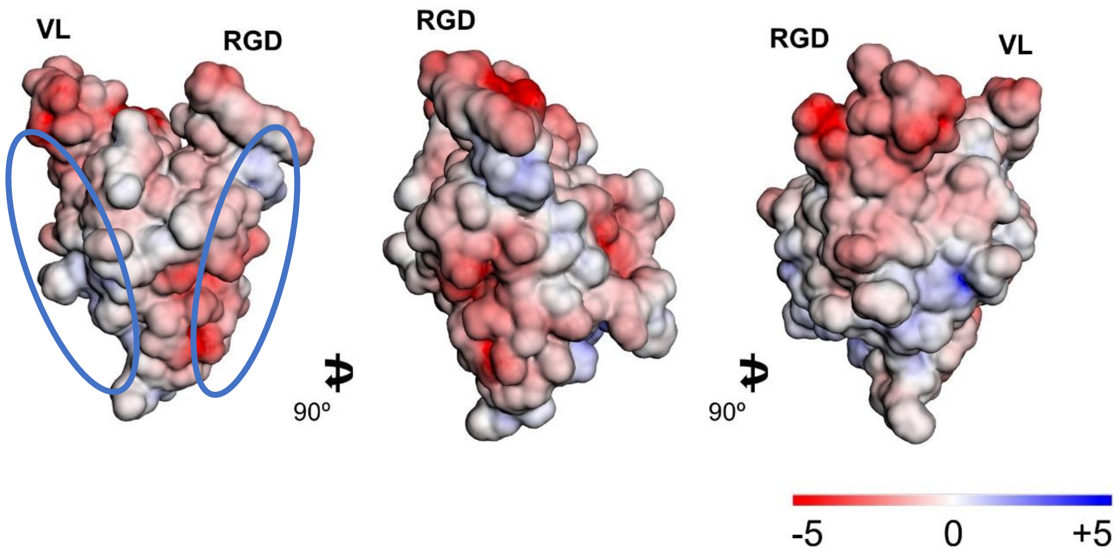


**Figure S3. Electrostatic (Coulomb) potential surface presentation of ADDobody** in a front, side and back view (scale in kcal mol $^{-1}$ e $^{-1}$) showing positively (blue) and negatively (red) charged patches. Interaction sites between ADDobodies forming a penton in the crystal are highlighted (blue circles). Loops (VL, RGD) are marked.

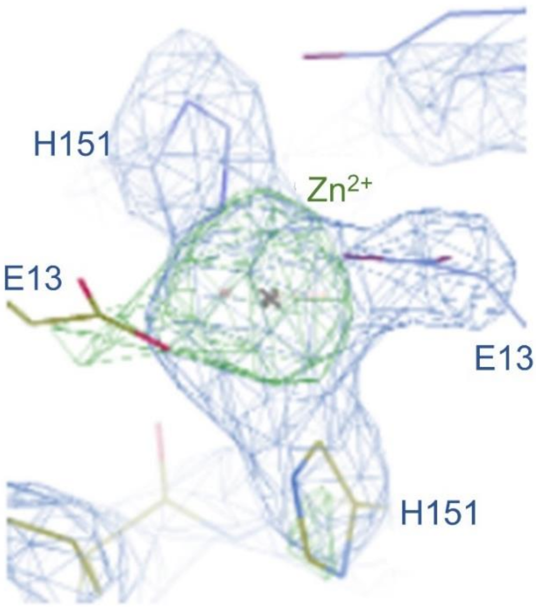


**Figure S4: Electron density for the coordinated $Zn^{2+}$ ion in the orthorhombic ADDobody crystal.** $Zn^{2+}$ binding stabilises the interaction between ADDobody penton rings in the crystal (see Fig. 2G). A close-up view of the electron density is shown at two different contour levels (0.5σ blue, 0.6σ green) for the $Zn^{2+}$ coordination by histidine H151 and glutamate E13 residues from ADDobodies juxtaposed in the lattice.

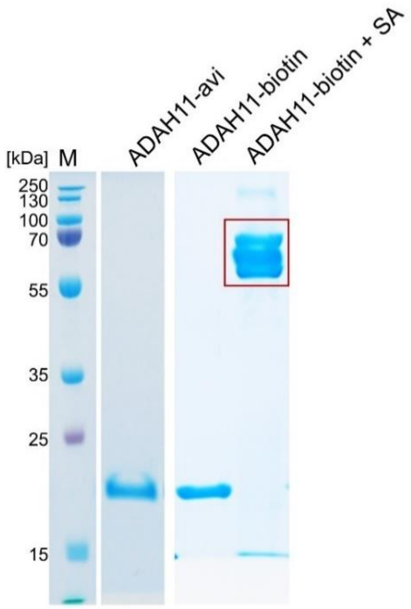


**Figure S5: ADAH11-avi purification and biotinylation.** Left: Coomassie-stained SDS gel section showing purified ADAH11 with a C-terminal avi tag (Molecular weight: 21.7 kDa). Right: gel shift assay using biotinylated ADAH11 and tetrameric streptavidin (SA, 52 kDa). Bands of the biotinylated ADDobody-Streptavidin complex run at higher molecular weight (red box) indicating close to quantitative biotinylation. M: Molecular weight marker.

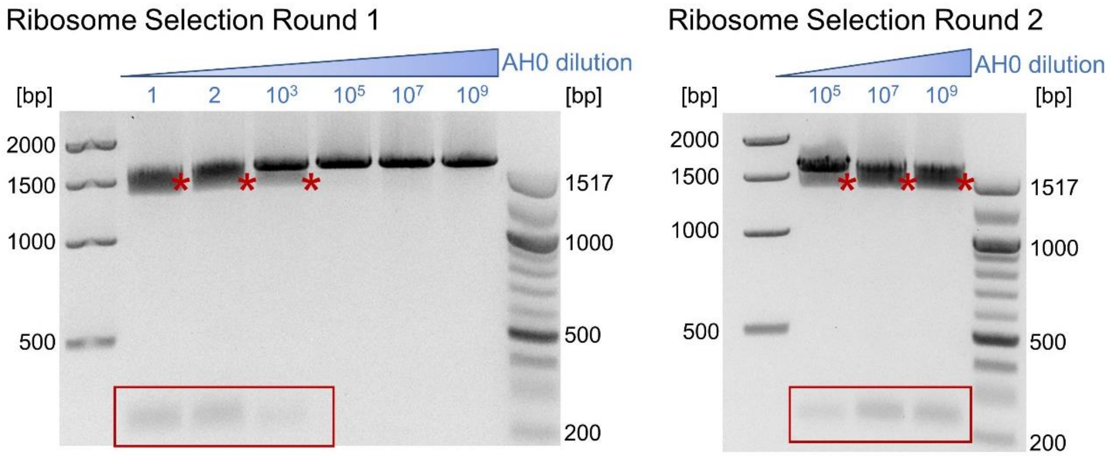


**Figure S6. Proof-of-concept ribosome display selections.** DNA agarose gel analysis of *Pst*I restriction digestions of the PCR products obtained after 1 and 2 rounds of ribosome display selection starting from different dilutions of ADDobody-AH0 into ADDobody-57 using ADAH11 as the target antigen. Numbers in blue indicate x-fold dilution. Bands corresponding to the cut PCR product (encoding ADDobody-AH0) are highlighted by a star (1,408 bp) and a box (242 bp).

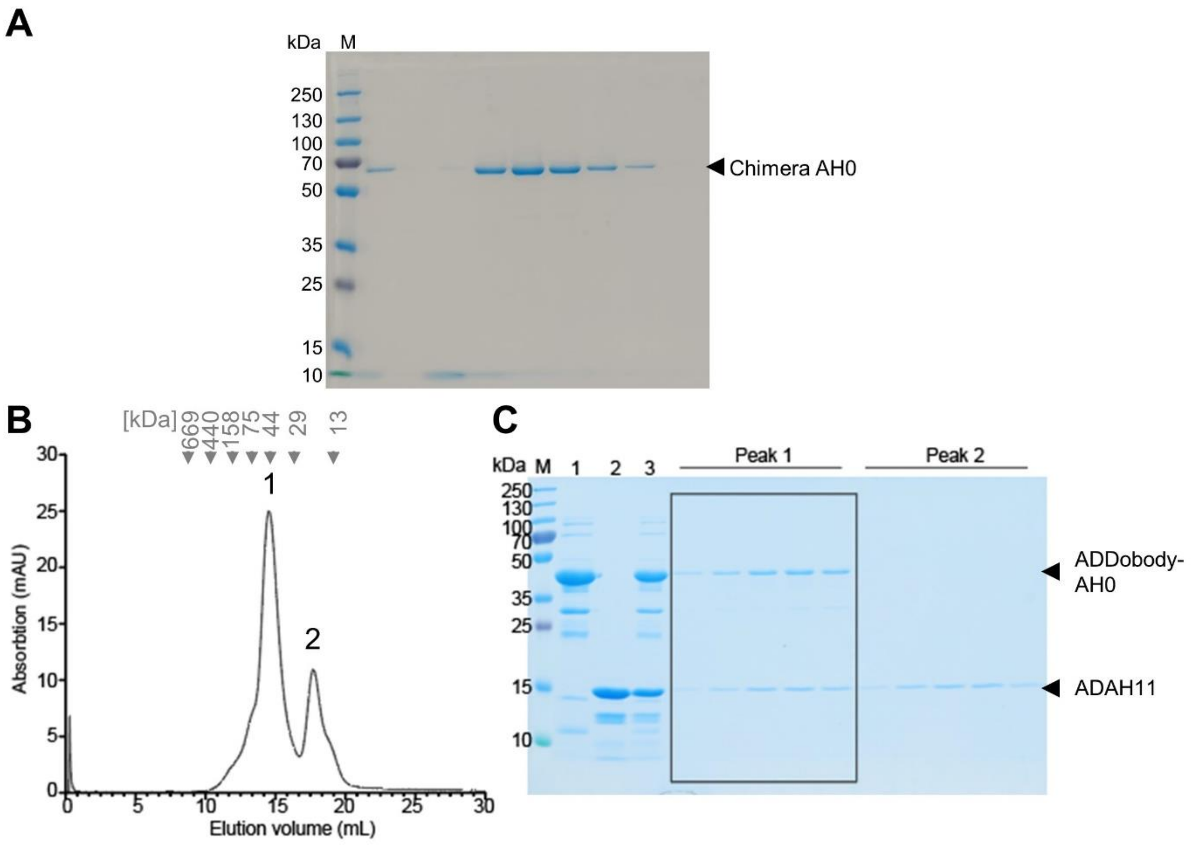


**Figure S7. Chimera AH0 purification and ADDobody-AH0 binding to ADAH11. (A)** Coomassie-stained SDS gel showing purified Chimera AH0 after ion exchange chromatography (IEX). The peak fraction of eluted protein was used for electron microscopy (Fig. S8). **(B)** Size-exclusion chromatography profile of a mix of ADDobody-AH0 and ADAH11 shows co-elution of the proteins, evidencing complex formation. ADDobody-AH0 and ADAH11 were mixed in a 1:1.5 molar ratio prior to loading onto a Superdex 200 10/300 GL column. **(C)** Coomassie-stained SDS gel showing the purified input samples ADDobody-AH0 (lane 1), ADAH11 target antigen (lane 2) and the mixture of the two proteins that was loaded on the column (lane 3). Fractions corresponding to Peak1 contain a complex between ADDobody-AH0 and ADAH11. Fractions corresponding to Peak2 contain unbound excess ADAH11. M: Molecular weight marker.

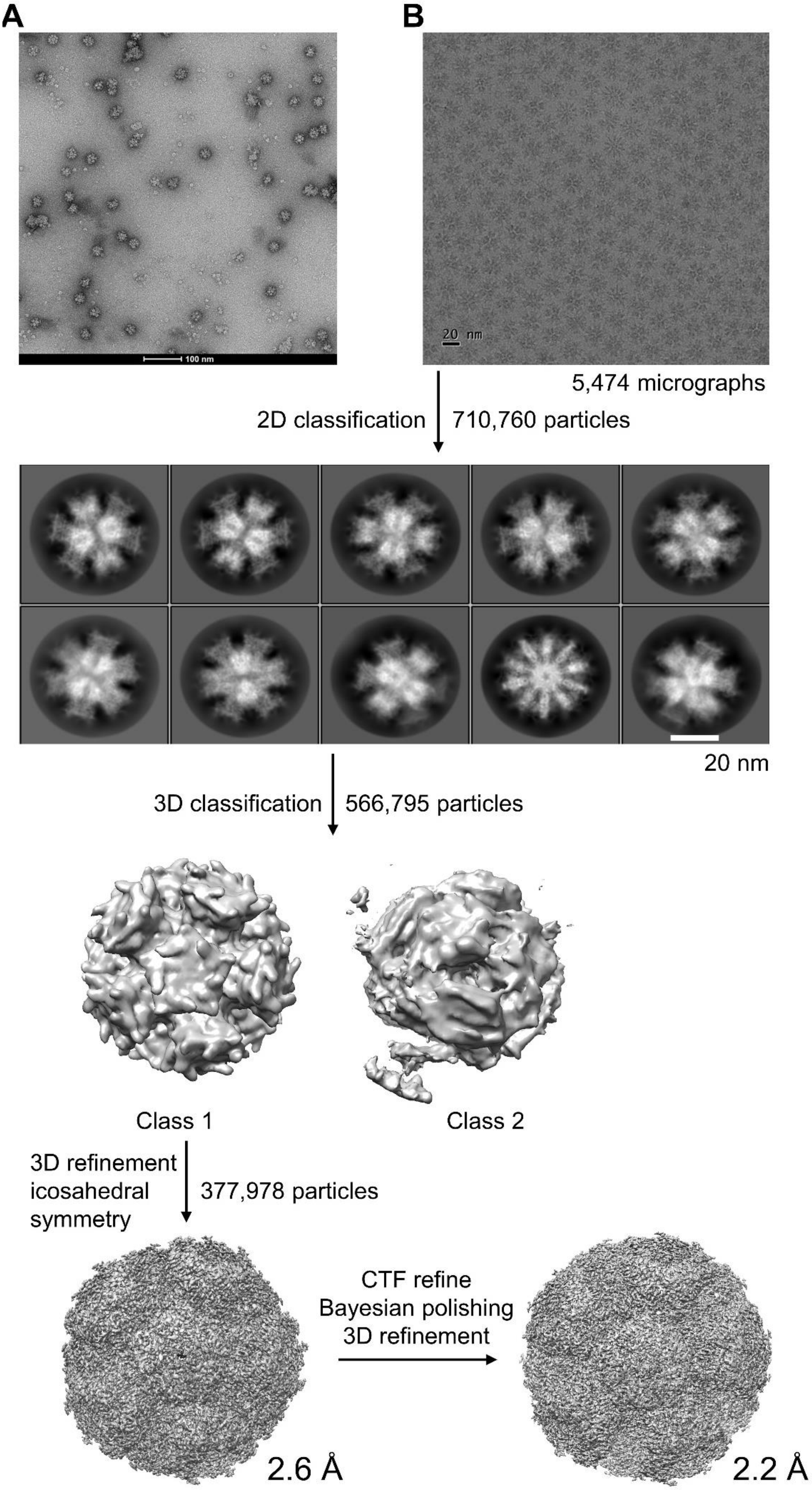
A
B
100 nm
20 nm
5,474 micrographs
2D classification
710,760 particles
20 nm
3D classification
566,795 particles
Class 1
Class 2
3D refinement
icosahedral
symmetry
377,978 particles
CTF refine
Bayesian polishing
3D refinement
2.6 Å
2.2 Å

**Figure S8. Electron microscopy of Chimera AH0. (A)** A representative negative stain EM micrograph of Chimera AH0. Scale bar: 100nm **(B)** Cryo-EM image processing workflow. A motion-corrected cryo-EM micrograph (scale bar 20 nm), reference-free 2D class averages (scale bar 20 nm), 3D classification, application of icosahedral symmetry and 3D refinement resulted in a 2.2 Å cryo-EM map.

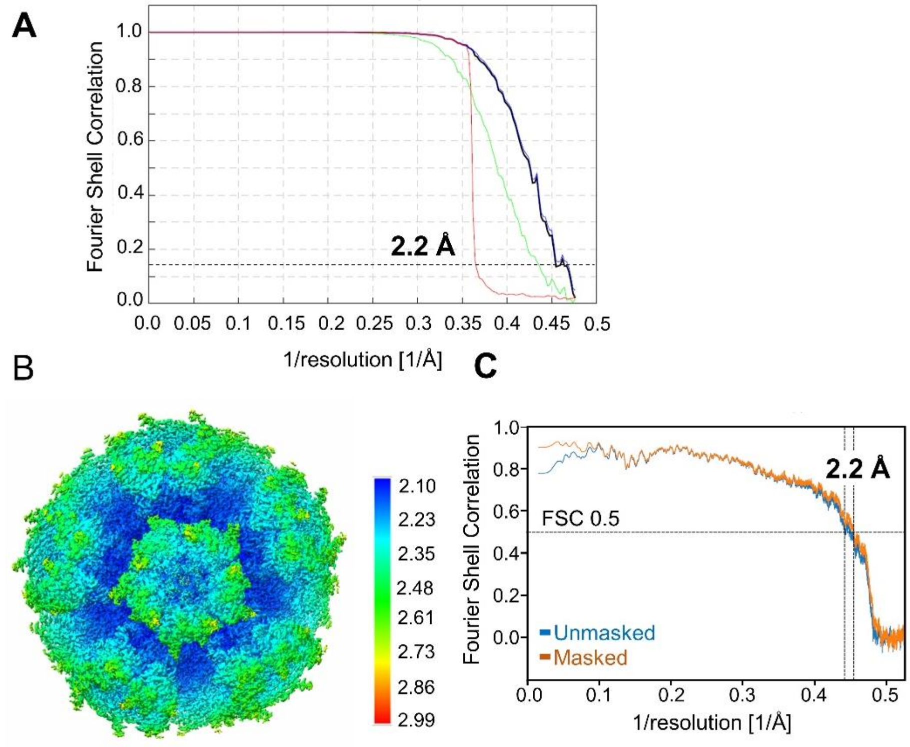


**Figure S9. Chimera cryo-EM map and model resolution**. **(A)** The Fourier Shell correlation (FSC) curve after gold-standard refinement of 377,978 particles. The FSC = 0.143 criterion indicates an overall resolution of 2.2 Å. Blue curve: FSC curve of masked map, green curve: FSC curve of unmasked maps; red curve: FSC curve of phase randomized masked maps. **(B)** Local resolution of the final Chimera cryo-EM map calculated in RELION 3.1. The core of the complex is resolved at 2.2 Å whereas peripheral parts comprising the VL and RGD loops have a lower resolution of ~ 2.7 Å. **(C)** FSC curve calculated between the atomic model and the final cryo-EM map. The map/model FSC at 0.5 reaches a resolution of 2.2 Å.

**Table S1: Amino-acid sequences of ADDobodies used in this study.**

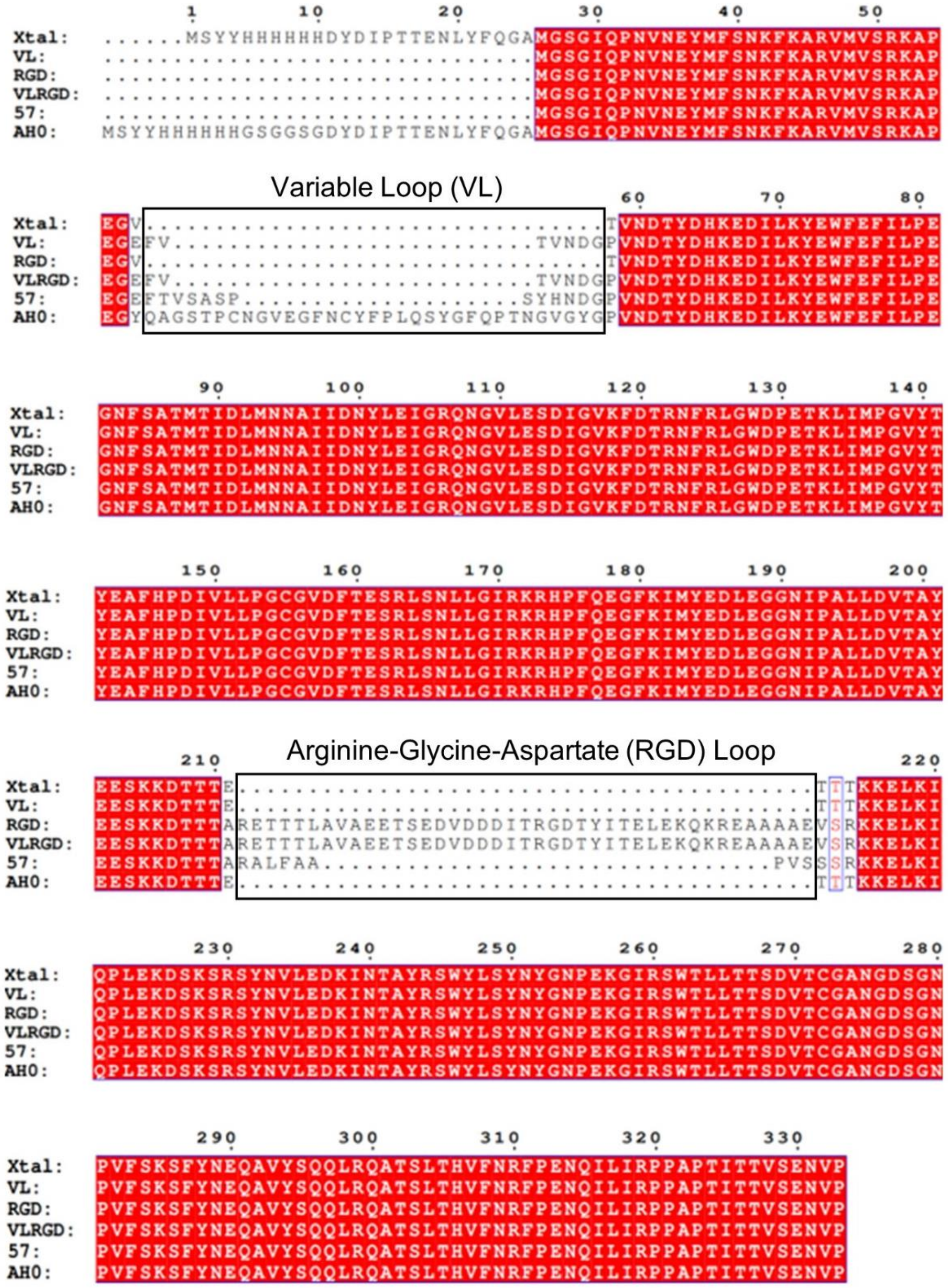

**Table S2. ADDobody crystallographic data collection and refinement statistics**

| **Data collection** | | |
|---|---|---|
| Structure | **zinc-free ADDobody** | **zinc-containing ADDobody** |
| PDB code | 8COI | 8QB3 |
| Wavelength (Å) | 0.9795 | 0.9780 |
| Synchrotron (Beamline) | Diamond Light Source (I03) | Diamond Light Source (I04) |
| **Cell Dimension** | | |
| Space group | P1 | P $2_1$ $2_1$ $2_1$ |
| a, b, c (Å) | 103.85 104.63 180.49 | 101.48 103.07 167.22 |
| α, β, γ (º) | 92.03 95.65 112.61 | 90.00 90.00 90.00 |
| **Collection Statistics** | | |
| Resolution range (Å) | 96.41 - 3.169 | 87.74 – 2.90 |
| Completeness (%) | 98.6 | 100 |
| Rmerge | 0.084 | 0.136 |
| Rmeas | 0.119 | 0.192 |
| CC 1/2 | 0.997 | 0.946 |
| Signal-to-Noise-Ratio (I/σ) | 1.44 | 1.06 |
| Total reflections | 420888 | 75083 |
| Unique reflections | 116474 | 39593 |
| Multiplicity | 3.6 | 1.9 |
| **Refinement Statistics** | | |
| Rwork (%) | 22.61 | 25.52 |
| Rfree (%) | 28.16 | 32.50 |
| RMS Bonds (Å) | 0.019 | 0.011 |
| RMS Angles (°) | 1.830 | 1.228 |
| Ramachandran Favoured (%) | 91.56 | 89.68 |
| Ramachandran Allowed (%) | 6.63 | 9.02 |
| Ramachandran Outliers (%) | 1.81 | 1.80 |
| Rotamer Outliers (%) | 2.7 | 2.50 |
| Wilson B-factor (Å$^2$) | 82.40 | 30.80 |
| Clash Score | 20.84 | 13.93 |
| MolProbity Score | 2.28 | 2.21 |

**Table S3. Cryo-EM data collection and refinement statistics, Chimera AH0**

| | |
|---|---|
| Voltage (kV) | 200 |
| Magnification (nominal) | 130,000 |
| Pixel size (Å/pix) | 1.05 (0.525) |
| Flux (e-/pix/s) | 5.9 |
| Frames per exposure | 40 |
| Exposure (e-/ Å2) | 1.06 |
| Defocus range (µm) | -0.8 to -2.0 |
| Micrographs collected | 6179 |
| Particles, final | 377,978 |
| Map sharpening B-factor (Å2) | -88 |
| Masked resolution at 0.143 FSC (Å) | 2.2 |

**Refinement**

| | |
|---|---|
| Composition | |
| Amino acids | 24180 |
| RMSD bonds (Å) | 0.004 |
| RMSD angles (°) | 0.747 |
| Mean B-factor (Å) | |
| Amino acids | 8.02 |
| Ramachandran | |
| Favoured (%) | 94.30 |
| Allowed (%) | 5.70 |
| Outliers (%) | 0 |
| Rotamer outlier (%) | 0.64 |
| Clash score | 1.84 |
| C-beta outliers (%) | 0 |
| CC (mask) | 0.80 |
| MolProbity score | 1.33 |
| EMRinger score | 5.83 |
| Model resolution (Å) 0.5 FSC threshold | 2.2 |

**Table S4. Ribosome display primers used in this study.**

| Primer | Sequence (5’ to 3’) |
|---|---|
| T7B_F_v3 | 5’ ATACGAAATTAATACGACTCACTATAGGGAGACCACAAC GGTTTCCCTCTAGAAATAATTTTG 3’ |
| A1.MS-SDA-ADDobody-F | 5’ AGACCACAACGGTTTCCCTCTAGAAATAATTTTGTTTAA CTTTAAGAAGGAGATATATATGGGATCCGGAATTCAACC 3’ |
| tonBtot_R | 5’CCGCACACCAGTAAGGTGTGCGGTCAGGATATTCACCA CAATCCC 3’ |
| A14-tonB-ADDobody-R | 5’ CCGCACACCAGTAAGGTGTGCGGTCAGGATATTCAC 3’ |
